# GlyTrait Brings Insights into Functional Glycosylation

**DOI:** 10.1101/2024.07.09.602632

**Authors:** Bin Fu, Guoli Wang, Chenxin Li, Yueyue Li, Xuejiao Liu, Ying Zhang, Haojie Lu

## Abstract

Glycomics research often grapples with the interpretability and biological relevance of glycomics data. Using glycosylation derived traits are promising methods for more in-depth biological insights, yet no such bioinformatic tool exists for such task. Here, we developed GlyTrait, a Python-based framework designed to enhance glycomics analysis through the innovative calculation and interpretation of derived traits from N-glycome data. GlyTrait automates the derivation of biologically significant traits, shifting focus from mere glycan abundances to functional glycan properties such as branching and fucosylation. GlyTrait extends the well-established nomenclatures and definitions of derived traits in the N-glycomics community, allowing for fast exploration and analysis of N-glycome data effortlessly. Furthermore, with the well-designed formula grammar, custom derived traits could be materialized without any knowledge of coding. Besides, a two-step post-filtering process reduces information redundancy, maintaining only the most informative traits. Finally, subsequent statistical and interpretable machine learning analysis provide robust insights into the glycosylation patterns associated with disease states. This comprehensive approach not only improves the statistical power and sensitivity compared to traditional methods, but also enhances the interpretability of glycomics data. GlyTrait’s efficacy is demonstrated through the re-analysis of published glycoengineered CHO cell lines and visceral leishmaniasis patient data, alongside a newly conducted pilot study for hepatocellular carcinoma (HCC) N-glycan biomarker discovery. We are confident in GlyTrait’s potential to become an indispensable tool for the glycomics community.

**Significance Statement:** Glycomics data is often challenging to interpret and analyze for biological relevance. GlyTrait, a Python-based tool, addresses this by automatically calculating biologically significant traits from N-glycome data, focusing on functional properties like glycan branching and fucosylation rather than mere abundance. This tool enhances data interpretability, reduces redundancy through post-filtering, and employs statistical methods to uncover glycosylation patterns linked to diseases. Demonstrated on various published datasets, as well as a newly conducted pilot study with 165 samples for hepatocellular carcinoma N-glycan biomarker discovery, GlyTrait proves to be a powerful tool. It improves the understanding of glycosylation changes in conditions like cancer, thereby benefiting the glycomics community.

## Introduction

With the development of precise medicine, glycomics is gradually gaining attention, along with other omics fields including genomics, transcriptomics, proteomics, lipidomics and metabolomics. N-glycosylation, as the most comprehensively studied type of glycosylation, has been revealed by numerous studies to have connections with rheumatic disease, inflammation, cancer progression, and aging, making it a promising diagnostic or prognostic biomarker(1), or a drug target(2).

On one hand, with the advancement of instrumentation and sample processing techniques, there has been a salient improvement in N-glycan identification and quantification, in terms of identification capacity, sample throughput, and structural elucidation depth(2, 3). On the other hand, various bioinformatics tools have been developed for more fast and more user-friendly processing and analyzing N-glycomics data(4). The functions of these tools include glycan identification (e.g., GlycoWorkbench(5), AssignMALDI(6), and GlycoNote(7)) as well as downstream analysis (e.g., glycowork(8) and GlyCompare(9)).

Bioinformatics advancement in proteomics have significantly contributed to resolving critical biological inquires, including differentiating distinct medical conditions, and elucidating underlying mechanisms. Traditional molecular comparative methods, which treat molecules as independent variable, are rational for proteome for the template-based regulatory mechanisms of proteins. However, glycosylation is a much more complex biological process orchestrated by numerous factors including glycosylation related enzymes, epigenetics, substrates, and environment(10), making direct comparison of individual glycans less biologically insightful. Besides, glycomics data usually suffers from sparsity and heterogeneity(11), hindering direct comparison of samples.

To address the above-mentioned issues, the glycomics community are trying to find its way better elucidating the intrinsic picture of N-glycosylation. One of the attempts is to calculate the substructures of glycans, exemplified by GlyCompare(9), which uses a well-designed algorithm to aggregate redundant substructures. Another direction is to calculate known functional glycan epitopes, also called glycan motifs, to spot biologically functional structural subunits of glycans. Glycowork is an intact Python framework for glycan motif analysis(8), and its recent update(12) introduced systematic ways of motif calculation and statistical analysis. Apart from that, several researchers incorporated glycosylation biosynthesis pathway to calculate a set of features reflecting certain kinds of glycosylation step(13–16). The new set of features are often called the “derived” traits, in contrast to glycan abundances, namely the “direct” traits. A derived trait reflects a characteristic of the entire glycome, rather than focusing on a single glycan. Compared to glycan abundance, derived traits are more biologically related to activities of specific enzymes in the glycosylation pathway and underlying genetic polymorphisms, making them beneficial for understanding the functional relevance of obtained results(3).

Although the derived trait methodology has been utilized by glycomics studies(2), several hard nuts are to be cracked. Firstly, up to now, derived traits calculation is mostly based on manually defined formulas, which depend on the identified glycan repertoire, preventing their generalization. Secondly, apart from the cumbersome calculation process *per se*, the calculated derived traits suffer from information redundancy. Many traits provide no meaningful information for the given glycan set, while others may highly correlate with each other, leading to redundant information. It is hard to decide which traits are important beforehand, while manual reduction of the trait dimension based on arbitrary rules afterwards turns out to be subjective and inefficient. Finally, although derived traits are more biologically relevant and thus more interpretable, each trait reflects only a restricted fraction of the glycosylation landscape, bringing challenges to integrating these insights comprehensively. All these factors render an automated tool for calculating and analyzing glycan derived traits imperative.

Herein, we developed GlyTrait, a Python-based tool for one-stop derived-trait-based analysis for N-glycome data. First, GlyTrait efficiently calculates hundreds of derived traits in an automated manner from diverse glycomics quantification data. After that, GlyTrait reduces redundant information by a sophisticated post-filtering phase, retaining only the most informative traits. In its final stage, GlyTrait harnesses both statistical methods and advanced interpretable machine learning techniques to unravel glycosylation patterns and their relevance to phenotypic characteristics. We demonstrated the advantages of GlyTrait in various glycomics application scenarios including guiding glycoengineering and providing insights into cancer development. First, a simple glycoengineered CHO dataset(17) was re-analyzed for demonstration of the trait calculation functionality. Subsequently, we carried out a pilot study for serum N-glycan biomarker discovery for patients with hepatocellular carcinoma (HCC). Finally, we demonstrated the capability of GlyTrait to analyze sialic acid linkages by re-evaluating a previously published dataset of serum N-glycome from patients with visceral leishmaniasis (VL). GlyTrait provides a convenient and user-friendly way to calculate N-glycan derived traits, along with the commonly used statistical analysis integrated. We believe GlyTrait will benefit the glycomics community in better elucidating glycan structural changes, especially in the clinical usage.

## Results

### The GlyTrait workflow

GlyTrait is a framework for analyzing N-glycomics data comprehensively and insightfully. The workflow includes three main steps (Supplementary Fig. 1). Firstly, derived traits that are more biologically relevant than glycan abundance are crafted in an exhaustive manner (involving the “preprocessing”, “formulas”, “calculating meta-properties”, and “trait calculation” modules). This step converts the raw glycan dimension into a new derived trait dimension without much loss of information. We demonstrated this step using a published N-glycosylation dataset on glycoengineered CHO cell lines. Secondly, the new derived trait dimension is pruned to reduce information redundancy and further boost variables’ interpretability (the “post-filtering” module). Finally, powered by robust statistical approaches and interpretable machine learning techniques, the refined dimension is scrutinized to unravel the association of glycosylation with research questions (the “statistical analysis” and “interpretable machine learning” modules). The functionalities of the second the third step were demonstrated using a pilot study for HCC glycosylation biomarker discovery, with 165 samples in total. Finally, with special emphasis on sialic acid linkages, we re-analyzed a published dataset of visceral leishmaniasis patients’ serum N-glycome, proving its potential in glycan isomer analysis.

### GlyTrait extracts derived traits automatically with high efficiency

The first problem we seek to work out is the calculation of derived traits in an automated manner. Instead of using concrete glycans directly, we designed the GlyTrait Formula grammar to use meta properties as building blocks (see Supplementary Note 1 for instruction for creating custom formulas). A meta property is a basic structural property of glycans (Fig. 1a), for example, “the glycan type”, “the number of antennas”, and “the number of core fucoses”, etc. Using meta properties, trait formulas can become concise, easy to read and write, and highly scalable (see Supplementary Table 1 and 2 for all GlyTrait formulas). For example, the formula of CFc, “the proportion of core-fucosylated glycans within complex ones”, is “CFc = [nFc > 0] // [type == ‘complex’]”. Internally, GlyTrait parses the formula strings and leverages the Python library Numpy and Pandas to build a meta-property table for ultra-fast vectorized derived trait calculation (see Methods for details), allowing more than one hundred derived traits to be calculated in seconds on a standard laptop for dozens of glycans and hundreds of samples. Compared to traditional methods, which require researchers to spend days on formula curation and coding, GlyTrait streamlines this process, making it quick and effortless.

**Fig. 1.**
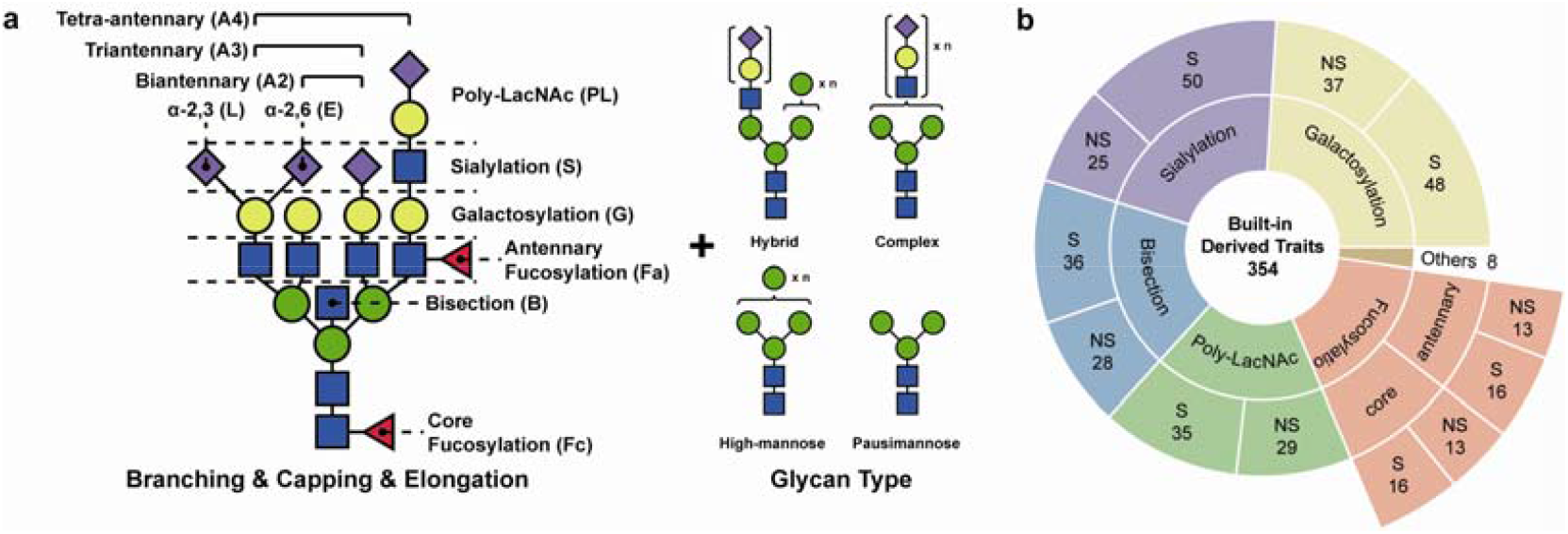
Overview of the GlyTrait framework for representing and calculating derived traits from N-glycome data. **a** The structural characteristics of N-glycans GlyTrait considers, including branching, common capping like fucosylation and sialylation, and glycan types, etc. **b** A summary of all 354 built-in derived traits supported by GlyTrait, classified into several categories and by whether sialic acid linkage is considered. “S” for “Sialic acid linkage considered”, “NS” for “Sialic acid linkage not considered”.

In total, 13 meta properties are defined by GlyTrait Formula grammar (Supplementary Note 1), constructing 354 built-in trait formulas (Fig. 1b, Supplementary Table 1), covering most appears in the highly-cited literature in a decade(13, 15, 16, 18–23), including traits about antenna number, glycan type, fucosylation, galactosylation, sialylation (with linkage information), bisection and poly-LacNAc elongation. Furthermore, powered by the well-defined GlyTrait formula grammar, GlyTrait supports custom trait formulas, making it a highly versatile and extensible tool. To maximize the usability, GlyTrait has utilities to help users rapidly prototype new formulas. GlyTrait can also run in the compositional mode when structural information is not available. In this mode, a set of different meta-properties and algorithms are used (Supplementary Table 2), deriving traits purely from glycan compositions. To ensure rigorous results, the compositional mode makes minimal assumptions about the structures. For example, the bisecting state will not be determined, and the locations of the fucoses will be ignored either. Finally, the users could always make their own structure annotations by providing an extra meta-property file to GlyTrait. This takes control of the meta-property extracting procedure from GlyTrait, combined with user-custom formulas, enable calculating derived traits for any types of glycans (O-glycans, milk oligosaccharides, or glycolipids). The latest documentation of GlyTrait could be accessed from https://github.com/FudanLuLab/glytrait.

We demonstrated the automated derived trait calculation functionality of GlyTrait using published erythropoietin (EPO) glycoprofiles from a series of glycoengineered CHO cell lines(17). Glycoengineering is crucial for the biopharmaceutical industry, as glycoprotein biologics is the fastest-growing class of therapeutics(24). In this dataset, 19 glycosyltransferase genes were knocked out individually or in a stacked manner to block certain glycosylation processes, resulting in diverse glycoprofiles. Direct qualitative determination of the effects of gene knockout could be achieved by comparing the most abundant glycan species in the spectra, as in the original paper. However, this approach lacks conciseness and intuitiveness. Also, quantitation of the knockout effects is impossible, for the sparsity of the glycans identified in different glycoprofiles prevented any useful pattern to emerge (Fig. 2a, Supplementary Data 1 – Input 1).

**Fig. 2.**
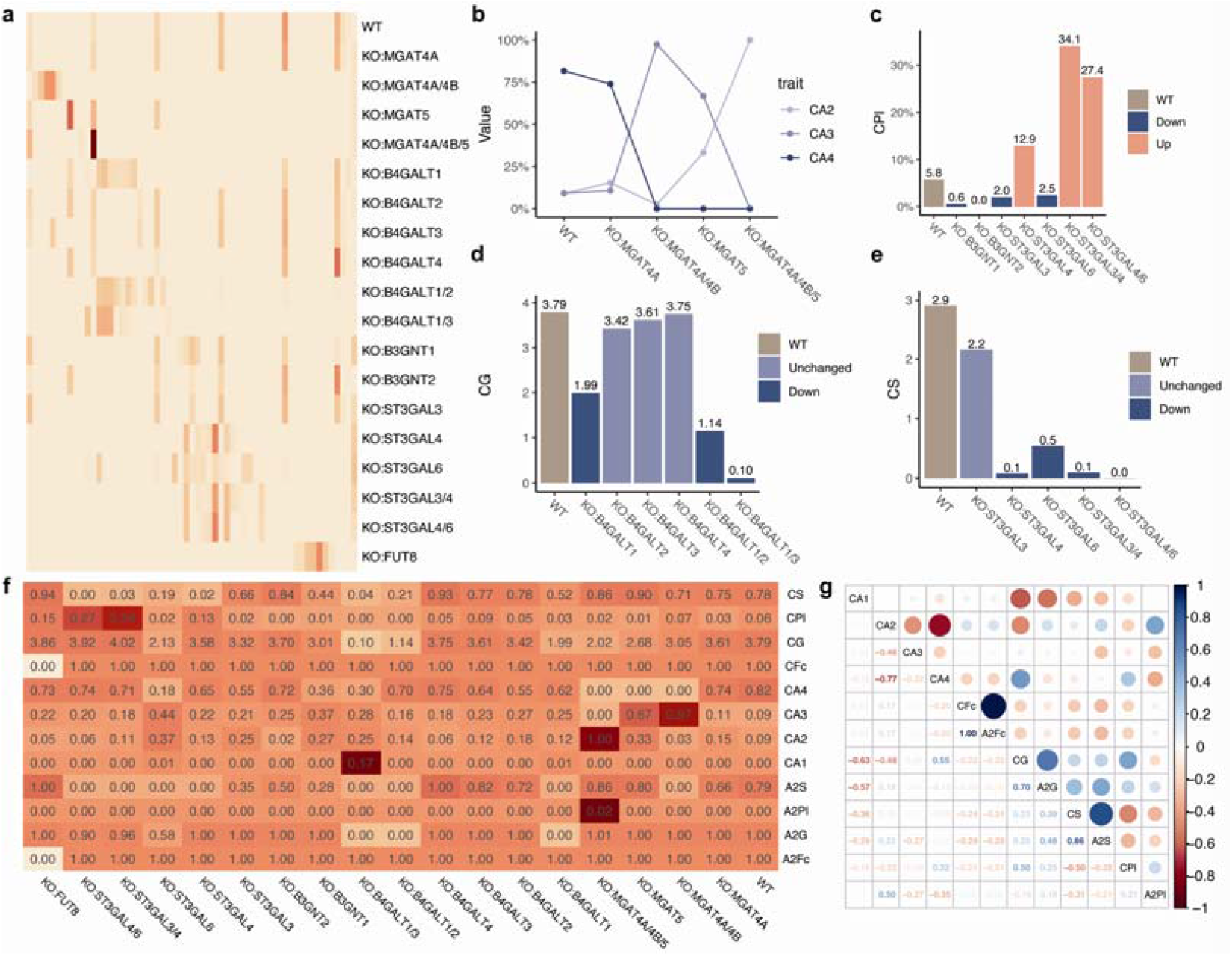
GlyTrait quantitively revealed N-glycosylation structure alteration for glycoengineered erythropoietins in CHO cell lines. **a** Glycoprofiles of the glycoengineered erythropoietins for different knockout samples showed sparsity and high heterogeneity, making direct comparison between samples not intuitive. KO: knockout, WT: wild type. **b** Relative abundance of glycans with different numbers of antennas (CA2, CA3, and CA4 for two, three and four antennas, respectively) in selected cell lines. **c-e** Relative abundance of glycans with (c) poly-LacNAc epitopes, (d) galactosylation, and (e) sialylation within glycans of complex type in selected cell lines. **f** Derived trait profiles of the dataset, with numbers in the cells being raw trait values. **g** The correlation relationships among derived traits implied certain degree of information redundancy (CFc vs A2Fc). Numbers in the cells are the Pearson’s coefficient of correlation.

We used GlyTrait to calculate all 152 built-in derived traits for the CHO datasets (Supplementary Data 1 – All derived traits). Compared to glycoprofiles, derived traits offer a more direct and quantitative view of glycosylation changes (Fig. 2b-f). To simplify the demonstration, only some representative derived traits are shown in Fig. 2f (see Table 1 for the definitions of the derived traits, and Supplementary Fig. 2 for the intact heatmap). GlyTrait integrally reproduced the conclusions in the original paper about branching (Fig. 2b), poly-LacNAc (Fig. 2c), galactosylation (Fig. 2d) and sialylation (Fig. 2e). Furthermore, by using derived traits, the effects of glycogene knockout can be quantified, and GlyTrait makes this process trivial. This underscores the tool’s ability to capture variations in glycosyltransferase activities, which is crucial for cohort glycomics studies.

**Table 1.**
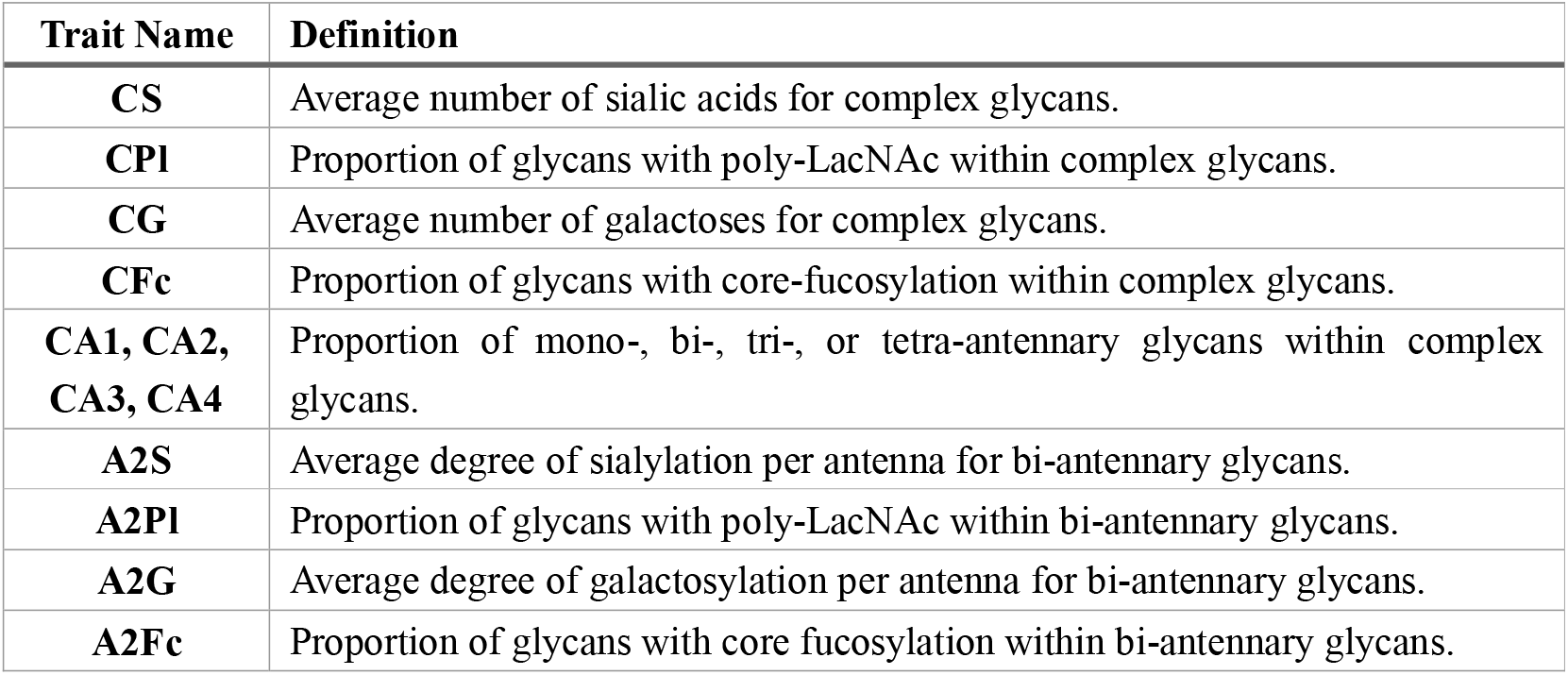
Definitions of selected derived traits in the CHO dataset. Only derived traits in Fig. 2 are explained here. For a comprehensive explanation of derived traits nomenclature, see Supplementary Table 1.

### Post-filtering reduces information redundancy in derived traits

During the above analysis, we found that most of the calculated derived traits for the CHO dataset were not informative. A case of point is the 28 bisecting traits, being zero in all samples (Supplementary Data 1 – All derived traits). This is because mannosyl (alpha-1,3-)-glycoprotein beta-1,2-N-acetylglucosaminyltransferase III (MGAT3), the glycosyltransferase adding bisecting N-acetylglucosamine to the pentasaccharide core, is not expressed in CHO cells. Also, some derived traits had large proportions of missing values (Supplementary Data 1 – All derived traits). Even among the remaining derived traits, a certain degree of information redundancy persisted (Fig. 2g, Supplementary Fig. 3). For example, the perfect collinearity between CFc and A2Fc (*r* = 1.00, *p* = 7.47×10^−166^) indicated identical extents of fucosylation within both biantennary complex glycans and all complex glycans. In this case, either CFc or A2Fc alone suffices to contain all the necessary information about the discrepancies among samples, making the other one redundant. These cases made the derived trait dimension highly non-informative and redundant, blurring the true underlying patterns of glycosylation alteration.

**Fig. 3.**
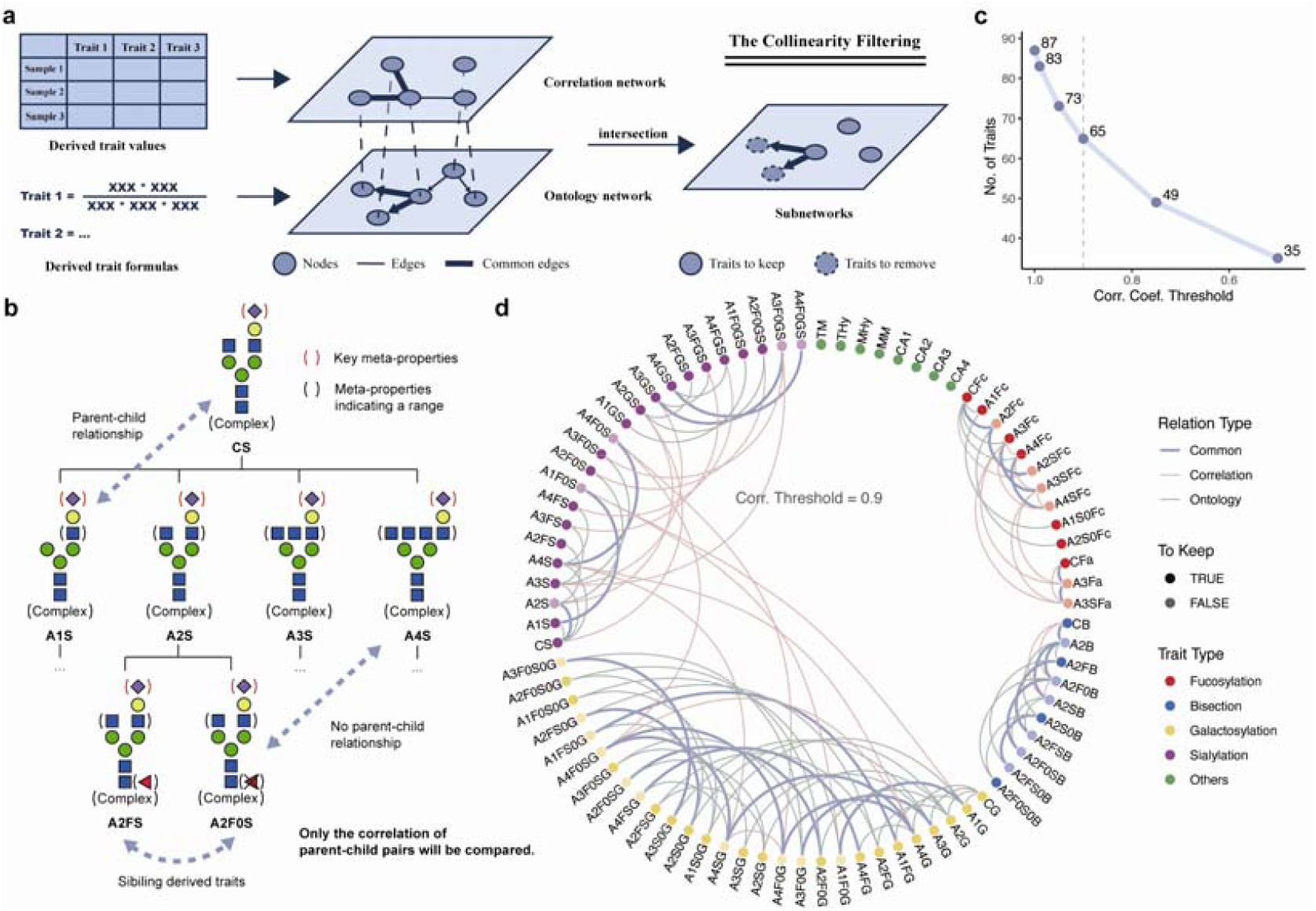
Post-filtering reduced information redundancy among N-glycan derived traits. **a** The collinearity filtering algorithm of GlyTrait. Briefly, a correlation network is built from the calculated derived trait values, and an ontology network from the definitions of the derived traits. The two networks are then merged into one, keeping only common edges in both networks. Only traits with zero in-degree in each component of the merged network are kept. **b** A part of the ontology network, focusing on derived traits about sialylation, with special focus on bi-antennary ones. In this zoomed part, CS is the parent of A1S, A2S, A3S and A4S, and A2S is the parent of A2FS and A2F0S. For the full network, see Supplementary Fig. 3, 4. **c** The number of retained derived traits after collinearity filtering with different correlation thresholds. **d** The relationships among derived traits, with correlation threshold of 0.9. The “common” edges connect traits with both high correlation and parent-child relationship. Darker dots represent derived traits retained in the collinearity filtering process.

To resolve this problem, we designed a post-filtering procedure consisting of two steps: one low-variance filtering step and one collinearity filtering step. The low-variance filtering simply removes derived traits with low variance and large proportions of missing values. The collinearity filtering algorithm is more complicated (Fig. 3a). Briefly, whenever two traits with hierarchical meanings highly correlated with each other (e.g. CFc and A2Fc), the one with a more general interpretation (CFc) will be kept, and the other (A2Fc) will be discarded (Fig. 3b, see Methods for more details). By this means, redundant traits will be removed, while potentially valuable correlation relationship will be retained. The filtering algorithm is tailored for derived traits, making the pruned trait dimension highly informative. Note that the correlation threshold of collinearity filtering could be set as a parameter of GlyTrait, allowing for tailored filtering for different research purposes.

To evaluate the efficiency of post-filtering in a more complex situation, we conducted a pilot study for depicting serum N-glycome alteration for hepatocellular carcinoma (HCC). One hundred and sixty-five serum samples were collected, including 55 HCC samples and 110 controls (54 hepatitis samples and 56 healthy controls). In total, 71 glycan compositions were detected by MALDI-TOF MS with structural annotations based on previous literature(25) (Supplementary Table 3, Supplementary Data 2 – Input 1 and Input 2), thus being used for further analysis. We then used GlyTrait to calculate all the 152 derived traits (Supplementary Data 2 – All derived traits). Ninety-one of them were kept after the low-variance filtering, and 65 of them (Supplementary Data 2 – Filtered derived traits; also see Table 2 for definitions of the derived traits) were kept after the collinearity filtering with a correlation coefficient threshold of 0.9 (Fig. 3c, d).

**Table 2.**
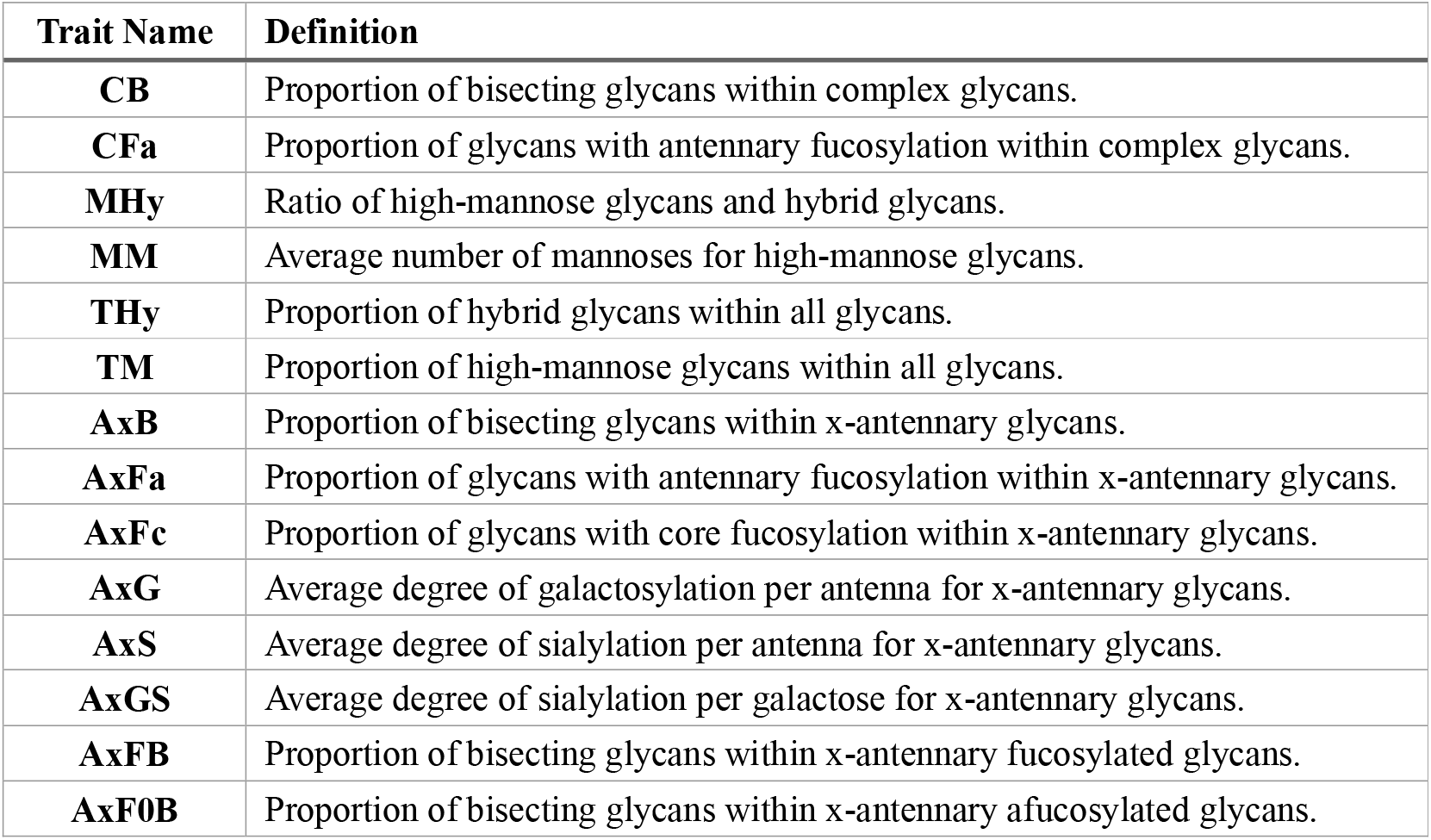
Definitions of selected derived traits in the HCC dataset. Derived traits already appears in Table 1 are skipped here. “x” can be either 1, 2, 3, or 4, denoting mono-, bi-, tri-, or tetra-antennary glycans. AxFB and AxF0B here are two examples of how middle qualifiers (F or F0) work. The meaning of other derived traits could be speculated following the same rule. For a comprehensive explanation of derived traits nomenclature, see Supplementary Table 1.

| **Trait Name** | **Definition** |
| --- | --- |
| **CB** | Proportion of bisecting glycans within complex glycans. |
| **CFa** | Proportion of glycans with antennary fucosylation within complex glycans. |
| **MHy** | Ratio of high-mannose glycans and hybrid glycans. |
| **MM** | Average number of mannoses for high-mannose glycans. |
| **THy** | Proportion of hybrid glycans within all glycans. |
| **TM** | Proportion of high-mannose glycans within all glycans. |
| **AxB** | Proportion of bisecting glycans within x-antennary glycans. |
| **AxFa** | Proportion of glycans with antennary fucosylation within x-antennary glycans. |
| **AxFc** | Proportion of glycans with core fucosylation within x-antennary glycans. |
| **AxG** | Average degree of galactosylation per antenna for x-antennary glycans. |
| **AxS** | Average degree of sialylation per antenna for x-antennary glycans. |
| **AxGS** | Average degree of sialylation per galactose for x-antennary glycans. |
| **AxFB** | Proportion of bisecting glycans within x-antennary fucosylated glycans. |
| **AxF0B** | Proportion of bisecting glycans within x-antennary afucosylated glycans. |

To examine whether the derived traits after post-filtering contain most information of the original glycan dimension, five widely used machine learning classification algorithms were trained on glycans-only and derived-traits-only features to predict sample group. Models were evaluated in 5-fold cross-validations, repeated for 10 times. The performance of most models showed no significant difference on two types of features, except for logistic regression (Supplementary Fig. 6). This proved that the transformation from glycan abundance to GlyTrait’s derived traits followed by post-filtering has negligible information loss.

### Statistical analysis by GlyTrait revealed biologically relevant glycosylation alteration

GlyTrait performed differential analysis after derived trait calculation and post-filtering when a file containing grouping information is passed in. By default, two-sided Welch *t*-test is used for two groups, and one-way Welch ANOVA with Games-Howell post-hoc test is used when more than two groups exist. Parametric tests were chosen over non-parametric ones considering the relatively larger sample sizes in modern glycomics studies (usually more than one hundred), where the central limit theorem (CLT) ensures that the sample means obey normal distributions regardless of the origin distribution of the population. However, it is well-known that diseases, especially cancers, manifest with a substantial degree of heterogeneity. To ensure robust statistical analysis, Welch correction of variance is performed for both *t*-test and ANOVA. All *p*-values are corrected for multiple comparison by Benjamini-Hochberg method. For *t*-test, Cohen’s *d* effect size is calculated, and partial *η*^2^is calculated for ANOVA.

To prove that derived-trait-based analysis has better statistical power than that by glycan-abundance-based analysis, we fabricated a simulated dataset (n = 100) using the 76 serum N-glycans with structure annotations(25). The 38 glycans with core fucosylation were scaled up to mimic the result of FUT8 up-regulation. When these glycans were scaled by 1.25 (simulating a fold change of 1.25 for each glycan), CFc showed statistical significance (*p* = 2.39×10^−5^), even after adjustment for multiple comparison with other derived traits (FDR = 2.27×10^−3^) (Supplementary Fig. 7). However, no glycans show statistical significance even before adjustment (Supplementary Fig. 8). When the scaling factor raised to 1.5, the up-regulation for only three of the 38 core-fucosylated glycans were spotted by *t*-test after adjustment (Supplementary Fig. 8), while CFc showed extremely confident significance (FDR = 2.19×10^−11^) (Supplementary Fig. 7). This underscores the statistical power and sensitivity of derived-trait-based analysis in detecting subtle biological variations that might be overlooked by traditional abundance-based approaches.

Armed with this boost in sensitivity and statistical power, we performed comparative glycomics analysis on the above-mentioned HCC dataset. Of the 65 derived traits kept by post-filtering, 36 were significantly different between HCC samples and controls, 21 of which increased in HCC samples and 15 decreased (Fig. 4a and Supplementary Fig. 9). The increased derived traits were miscellaneous regarding to fucosylation, sialylation, and bisection, while the decreased ones were mainly about galactosylation. Fig. 4b summarizes the alteration of 13 most fundamental and general derived traits. While mono-antennary glycans remained unchanged (CA1, FDR = 0.972), bi-antennary glycans (CA2, FDR = 1.33×10^−4^, Cohen’s *d* = −0.80) decreased, and tri-antennary (CA3, FDR = 5.34×10^−4^, Cohen’s *d* = 0.72) and tetra-antennary glycans (CA4, FDR = 6.16×10^−6^, Cohen’s *d* = 1.00) increased. Notably, core fucosylation level (CFc, FDR = 3.23×10^−7^, Cohen’s *d* = 1.09) increased, while antennary fucosylation (CFa, FDR = 0.170) remained stable. The proportion of hybrid glycans (Thy, FDR = 4.50×10^−3^, Cohen’s *d* = 0.53) increased, while the ratio of high-mannose glycans and hybrid glycans (MHy, FDR = 5.80×10^−8^, Cohen’s *d* = −1.00) decreased. Finally, bisection increased in HCC (CB, FDR = 5.37×10^−4^, Cohen’s *d* = 0.71).

**Fig. 4.**
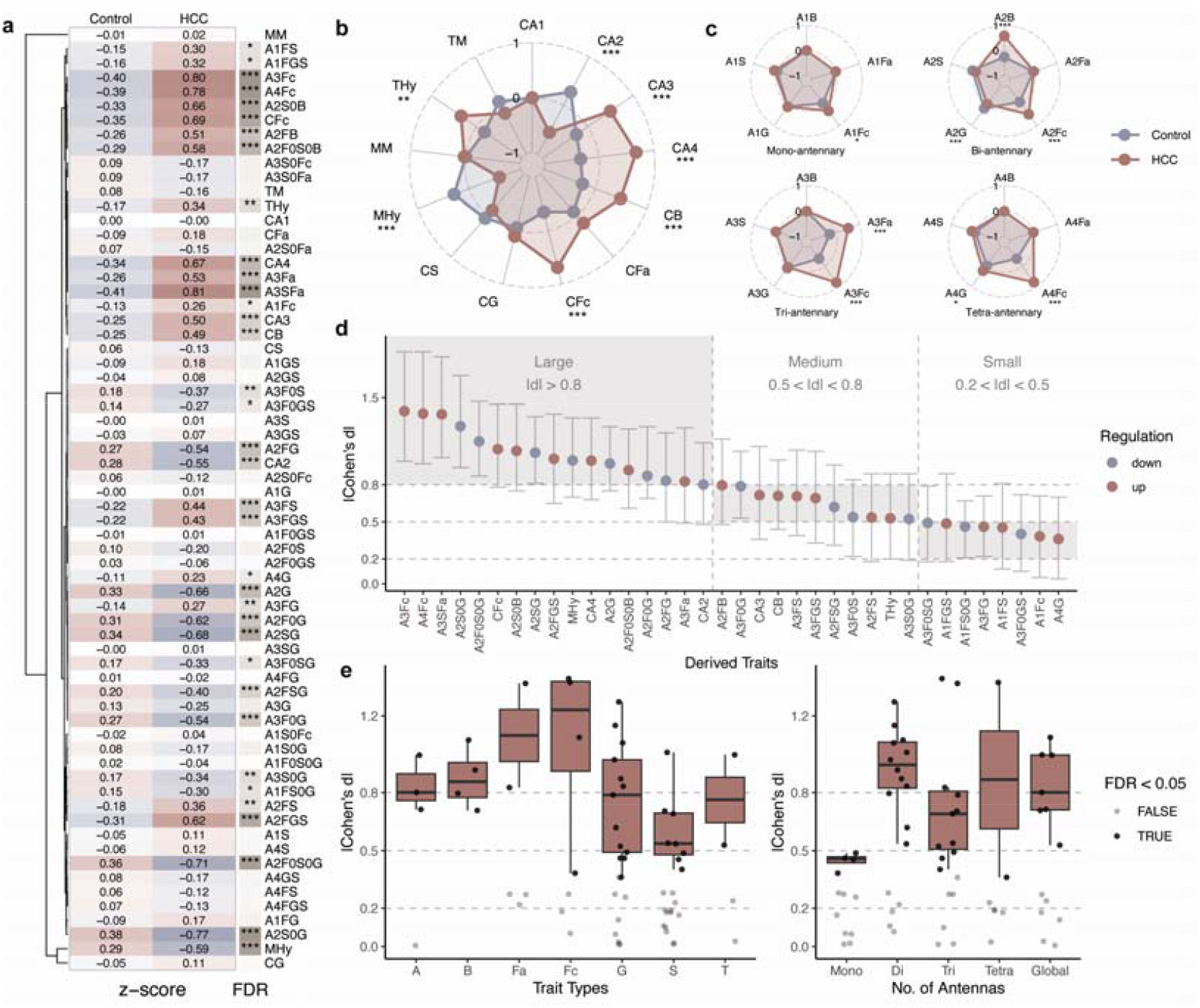
Glycosylation alteration in HCC serum samples was revealed comprehensively by statistical analyzing approach of GlyTrait. **a** 35 of 65 derived traits after post-filtering showed significant difference in Welch *t*-test (FDR < 0.05) between HCC and control samples. Numbers in the cells are average trait values for each group after z-score normalization to all samples. FDR: false discovery rate. ^*^: 0.01 < FDR < 0.05, ^**^: 0.001 < FDR < 0.01, ^***^: FDR < 0.001. The notation is consistent for figures afterwards. **b** Comparison between HCC and control samples for 13 global derived traits. Derived trait values were scaled into z-scores. **c** Comparison of core fucosylation (Fc), antennary fucosylation (Fa), bisection (B), galactosylation (G), and sialylation (S) for glycans with different numbers of antennas. **d** Effect sizes (Cohen’s d) for significant derived traits. J. Cohen’s criteria were adopted. Error bars show 95% CI. **e** The distribution of effect sizes (Cohen’s d) for different types of derived traits (left), and derived traits of different number of antennas (right). Boxplots are based only on significant derived traits. A: branching, B: bisection, Fa: antennary fucosylation, Fc: core fucosylation, G: galactosylation, S: sialylation, T: glycan types.

When scrutinizing the glycosylation properties for glycans with different numbers of antennas, more patterns surfaced (Fig. 4c). Core fucosylation level elevated in glycans with any number of antennas, while the amplitude was tremendously larger in tri- and tetra-antennary glycans (A3Fc, FDR = 9.67×10^−11^, Cohen’s *d* = 1.39; A4Fc, FDR = 8.32×10^−11^, Cohen’s *d* = 1.37). In fact, these two traits were the top two with largest effect sizes (Fig 4d). Interestingly, antennary fucosylation only increased in tri-antennary glycans (A3Fa, FDR = 1.90×10^−5^, Cohen’s *d* = 0.83). This might be due to the relative low detection rates for glycans with antennary fucosylation in serum (only H4N4F3S1, H5N4F2, H5N5F2 and H6N5F2S3). Similarly, bisection only increased in bi-antennary glycans, for only bi-antennary glycans with bisection were confidently identified. Finally, although not differed much in fold changes, galactosylation in bi-antennary glycans decreased with a large effect size (A2G, FDR = 7.87×10^−6^, Cohen’s *d* = −0.97).

Of the 36 significant derived traits, 17 had large effect sizes (|Cohen’s *d*| > 0.8), 11 had medium effect sizes (0.5 < |Cohen’s *d*| < 0.8), and 8 had small effect sizes (0.2 < |Cohen’s *d*| < 0.5) (Fig. 4d). Six of the derived traits with large effect sizes were about galactosylation in bi-antennary glycans, which all showed decreasing trends. Besides, 3 derived traits about core fucosylation (A3Fc, A4Fc, CFc) ranked top. A3SFa (FDR = 9.17×10^−10^, Cohen’s *d* = 1.37), A2S0B (FDR = 1.33×10^−7^, Cohen’s *d* = 1.07), and A2FGS (FDR = 3.49×10^−7^, Cohen’s *d* = 1.01) also ranked top. The 65 derived traits were then classified into 7 categories (Fig. 4e, left). Galactosylation traits and fucosylation traits turned out to have larger effect sizes, while sialylation traits had relatively lower effect sizes. When the derived traits were classified based on the number of antennas, bi-antennary traits appeared to have larger effect sizes, while mono-antennary traits had the least effect sizes (Fig. 4e, right).

Finally, we evaluated the potentiality for these derived traits to be novel biomarkers of HCC using ROC analysis with bootstrap (n = 1000) (Supplementary Fig. 10). Seven derived traits had an ROC AUC above 0.8, including A4Fc (AUC [95% CI] = 0.834 [0.758, 0.901]), A3Fc (0.831 [0.758, 0.897]), A3SFa (0.831 [0.754, 0.895]), A2S0G (0.819 [0.749, 0.883]), A2F0S0G (0.814 [0.740, 0.880]), A2F0G (0.813 [0.738, 0.888]), and A2SG (0.807 [0.731, 0.872]).

### Interpretable machine learning revealed key glycosylation traits in cancer development

Statistical methods provided a detailed view of the glycosylation alteration from control to HCC samples. However, they assumed a linear relationship between features and targets, and failed to capture the interactions among features. Furthermore, when derived traits were grouped into categories, it was not easy to aggregate the results. Thereafter, we turned to explainable machine learning (ML) techniques and integrated it into the GlyTrait workflow. Briefly, we use XGBoost trained on derived traits, combined with SHapley Additive exPlanations (SHAP), to provide a comprehensive understanding of glycosylation pattern alteration in HCC samples (Fig. 5a). XGBoost was chosen for its superior performance in tabular data, even better than deep learning in many situations(26). Feature importance is acquired by calculating the average absolute Shapley values among all samples, which represents the average degree of contribution of the trait to the predicted probability (denoted by mean |ΔP|). For a robust importance evaluation, the significance of each feature importance score is acquired from null models generated from 1000 permutations (denoted by *p*_*perm*_, see Methods).

**Fig. 5.**
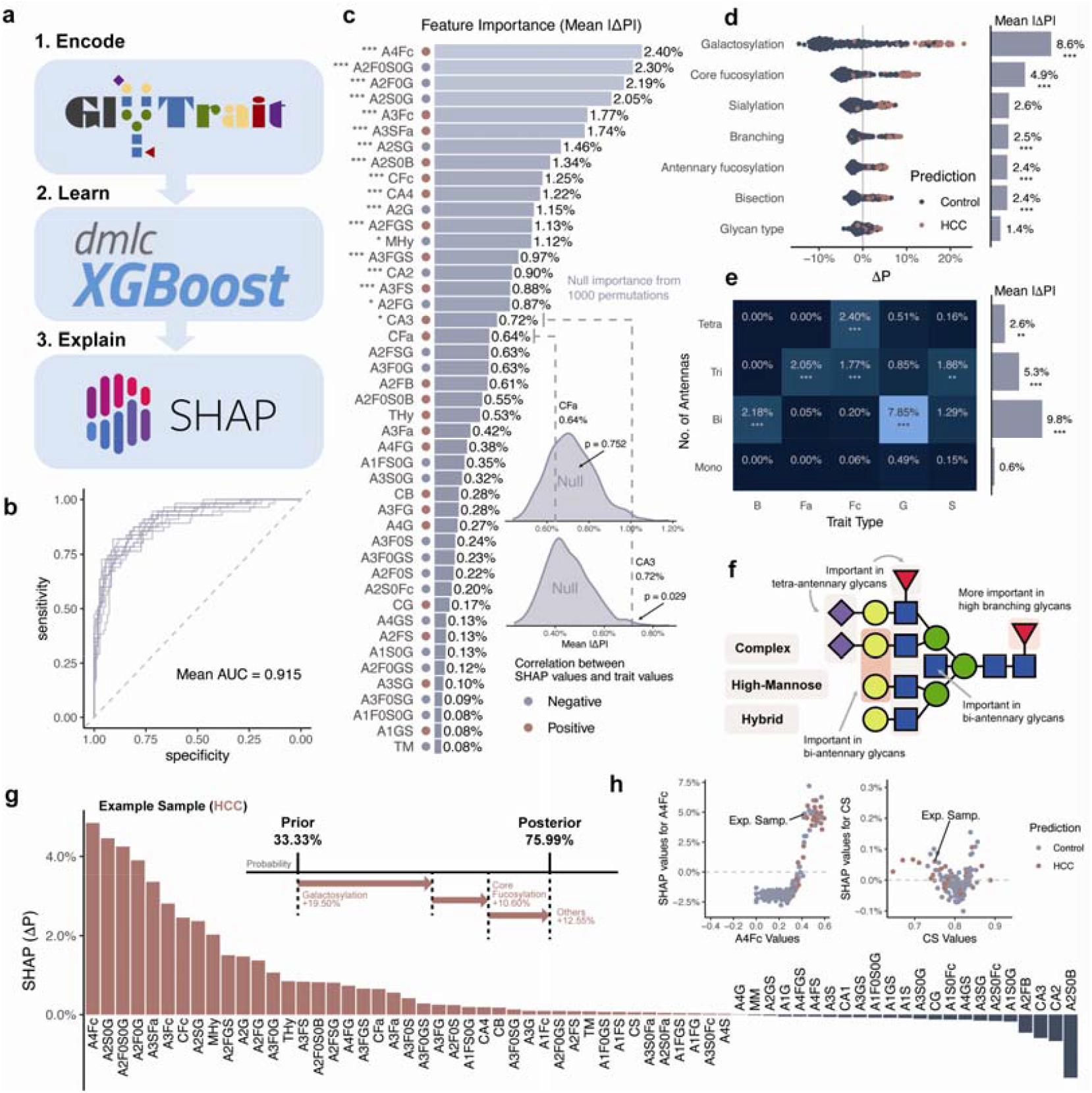
GlyTrait leverages interpretable machine learning techniques to mine the patterns in the glycomics data, exemplified on the HCC serum dataset. **a** The brief workflow of the machine learning strategy of GlyTrait. The model fine-tune, model evaluation, and Shapley values calculation are executed in a nested cross validation manner. **b** ROC plots for the fine-tuned XGBoost models evaluated in all outer layer repeated cross validation splits. Each line is a ROC curve using all predicted probability in cross validation in a repeated split. **c** The importance of the 65 derived traits after post-filtering given by SHapley Additive exPlanations (SHAP) algorithm. Importance of a trait is calculated as the average absolute Shapley values of the trait across all samples. Significance of the importance is denoted as stars: ^*^: 0.01 < FDR < 0.05, ^**^: 0.001 < FDR < 0.01, ^***^: FDR < 0.001. The contributing direction of a derived trait could be acquired by the correlation relationship of the trait values and the Shapley values (the red and blue dots), or the SHAP dependency plots (Supplementary Fig. 9). **d** SHAP summary plot of derived trait groups. A wider dispersion of samples implies a larger importance. **e** Importance of more detailed trait groups considering number of antennas. B: bisection, Fa: antennary fucosylation, Fc: core fucosylation, G: galactosylation, S: sialylation. **f** A summary of the importance of various glycosylation properties. **g** The Shapley values of an exemplified HCC sample (bottom) and a SHAP force plot for the explanation of model prediction for this sample (top). The selected sample had an average predicted probability of 75.99% in 10 repeated cross validation splits. The Shapley values were calculated in the probability space, so each value is the contribution of a trait towards the predicted probability. **h** The SHAP dependency plots for A4Fc and CS, as the examples for an important trait and an unimportant one, respectively.

The XGBoost model showed decent performance across all repeated splits, with an average ROC AUC of 0.915 ± 0.048 (Fig. 5b) on our dataset, slightly better than the random forest models before. The performance of the fine-tuned models on the training sets (AUC = 0.870 ± 0.094) was similar with that on the test sets, proving no apparent overfitting, thus the models could reflect the underlying data pattern authentically.

Of all 65 derived traits used for training the models, 19 had significant importance (*p*_*perm*_ < 0.05, after Benjamini-Hochberg adjustment) (Fig. 5c). Of the 19 derived traits, 6 were about galactosylation in bi-antennary glycans (A2F0S0G, A2F0G, A2S0G, A2SG, A2G, A2FG), 4 were about core fucosylation (A4Fc, A3Fc, CFc, A3S0Fc; A3S0Fc not shown in Fig. 5c for low importance), 3 were about sialylation in fucosylated glycans (A2FGS, A3FGS, A3FS).

Shapley values are additive, meaning that the contribution of a feature group in a prediction could be derived by calculating the summation of all Shapley values of those features, so the importance of a feature group could then be derived by calculating the mean absolute Shapley values for feature in the group. This property makes it possible to aggregate similar derived traits (Fig. 5d, e), which could not be easily achieved by statistical approaches. The galactosylation levels outstand the most (mean |ΔP| = 8.63±5.29%, *p*_*perm*_ < 0.001), followed by the core fucosylation levels (4.89±2.97%, *p*_*perm*_ < 0.001) (Fig. 5d). Similarly, derived traits could be aggregated by number of antennas (Fig. 5e, bar plot). Bi-antennary traits turned out to be most important (9.81±6.02%, *p*_*perm*_ < 0.001), followed by tri-antennary (5.34±3.16%, *p*_*perm*_ < 0.001) and tetra-antennary (2.59±1.67%, *p*_*perm*_ = 2.67×10^−3^) ones. When we further combined these two classification criteria, more patterns emerged (Fig. 5e, heatmap). Notably, galactosylation levels of bi-antennary glycans contributed largely (7.85±4.87%, *p*_*perm*_ < 0.001), shadowing all other feature groups. The core fucosylation levels turned out to be significantly important in high-branching glycans (tri-antennary, 1.77±1.11%, *p*_*perm*_ < 0.001; tetra-antennary, 2.40±1.45%, *p*_*perm*_ < 0.001). Furthermore, bisection was only vital in bi-antennary glycans (2.18±1.16%, *p*_*perm*_ < 0.001), and sialylation was only influential in tri-antennary glycans (1.86±0.87%, *p*_*perm*_ = 3.33×10^−3^). These results reveal the significance of glycan traits in predicting model outcomes, particularly highlighting the importance of galactosylation in bi-antennary glycans and core fucosylation in high-branching glycans (Fig. 5f).

Finally, it is worth mentioning that Shapley values could also be used to explain individual predictions of ML models. A Shapley value of a trait in a sample is the marginal contribution of this trait as a delta probability. One example was shown in Fig. 5g and 5h. Adding all ΔP values in the sample pushed the probability from 33.33% (the prior probability, i.e. the proportion of HCC samples within all samples) to 75.99%.

### GlyTrait provides insights into sialic acid linkages

Sialic acids are monosaccharides as part of many terminal glycan motifs of glycoconjugates found in vertebrates, involving in numerous biological processes(27). Studies have proven that sialic acid capping with different linkages differs in biological functions. With the advance of sialic acid linkage specific derivation techniques, these techniques have gradually been adopted for a better structural resolution. Sialylation on galactoses is catalyzed by two series of enzymes in mammals. ST6GAL family catalyzes α2,6-linked sialylation, while ST3GAL family catalyzes α2,3-linked sialylation.

To demonstrate the ability of GlyTrait for analyzing sialylation linkage, we re-analyzed a published dataset of total serum N-glycosylation from visceral leishmaniasis patients and controls(28). As the original article had depicted the glycosylation alteration comprehensively, here we only focus on the sialic acid linkage properties. Twenty-five derived traits about sialic acids linkages were kept after post-filtering (Fig. 6a). Among these, 20 differed significantly between visceral leishmaniasis (VL) and asymptomatic controls (Asy), 22 differed significantly between VL and healthy controls from endemic areas (EC). While the overall sialylation degree remained consistent for glycans of different numbers of antennas, we found a drop of α2,6-linked sialylation and an increase of α2,3-linked sialylation as the number of antennas increases (Fig. 6b). Overall, α2,6-linked sialylation decreased significantly (FDR = 1.58×10^−13^ between VL and Asy, FDR = 3.23×10^−2^ between VL and EC), which is consistent with the original paper. However, GlyTrait allowed for more detailed discoveries failed to be spotted before. Interestingly, we found that the degree of alteration (Cohen’s *d*) for α2,3-linked sialylation strongly correlated with numbers of antennas (Fig. 6c), increasing in high-branching glycans while decreasing in low-branching ones. This effect was not observed for α2,6-linked sialylation due to the obvious violation of bi-antennary glycans (Fig. 6d).

When scrutinizing sialylation degrees under the context of fucosylation, more patterns emerged (Fig. 6e-h). First, α2,6-linked sialylation degree increased for nearly all glycans regardless of fucosylation, with an exception for bi-antennary fucosylated glycans (A2FE: Cohen’s *d* = −1.06, FDR = 2.40×10^−24^ for VL vs. Asy; Cohen’s *d* = −0.72, FDR = 1.46×10^−10^ for VL vs. EC) (Fig 6e, 6g). Considering the moderate degree of core fucosylation within bi-antennary glycans (A2Fc, 28.1±8.8%), the down-regulation of α2,6-linked sialylation for bi-antennary glycans (Fig. 6d) might be due to this abnormality for bi-antennary fucosylated glycans. As for α2,3-linked sialylation (Fig. 6f, 6h), the decrease in low-branching glycans exclusively happened on afucosylated glycans, and the increase in tri-antennary glycans only existed for fucosylated glycans.

**Fig. 6.**
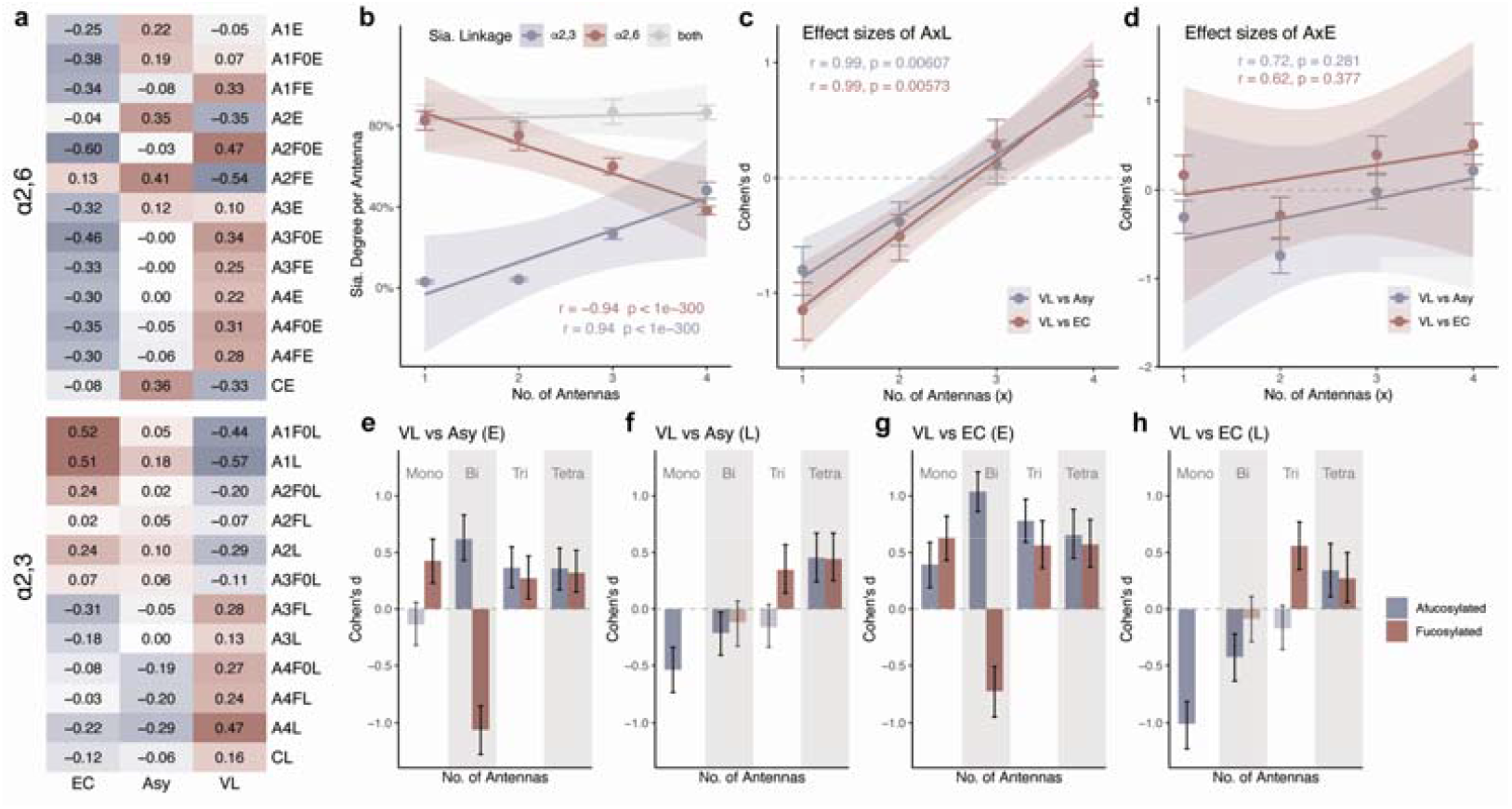
Derived traits about sialic acid linkages revealed key sialylation alteration in visceral leishmaniasis patients. **a** Overview of derived traits about sialic acid linkages. Numbers in the cells are average z-scores of trait values. EC: controls from endemic areas, Asy: ss. **b** Average degree of sialylation (average number of sialic acids per antenna) for glycans with different numbers of antennas. Error bars: standard errors of trait values. Shade: 95% CI of the linear models. **c-d** Effect sizes of AxL (c) and AxE (d) traits (x: the number of antennas, 1-4). AxL: average degree of α2,3-linked sialylation for glycans with x antennas. AxE: average degree of α2,6-linked sialylation for glycans with x antennas. Error bars: 95% CI of the effect sizes. Shade: 95% CI of the linear models. **e-h** The alteration of sialylation degree for glycans with (red) and without (blue) fucosylation, and with different numbers of antennas. e, g: α2,6-linked sialylation. f, h: α2,3-linked sialylation. e, f: Comparison between VL and Asy. g, h: Comparison between VL and EC. Error bars: 95% CI of the effect sizes. Bars with lighter color denote statistical insignificance. E: α2,6-linked sialylation, L: α2,3-linked sialylation.

## Discussion

Here we developed a comprehensive workflow for derived-trait-based comparative N-glycomics analysis and encapsulated it into a Python tool: GlyTrait. GlyTrait efficiently calculates derived traits in an exhaustive and automated manner, followed by a post-filtering step to reduce information redundancy. Finally, robust statistical analysis and interpretable machine learning techniques are integrated to find the association of N-glycosylation with specific research questions.

We demonstrated that the workflow yielded more biologically relevant results compared to traditional glycan-abundance-based analysis, with more interpretability and sensitivity. By re-analyzing the CHO dataset(17), similar conclusions were drawn from GlyTrait results as the original paper, proving the accuracy and validity of GlyTrait. Additionally, the alteration of glycosylation could be directly quantified by GlyTrait, which is lacking in the original paper. To demonstrate its potentiality in analyzing more complex dataset, we carried out a pilot study of serum N-glycan biomarker discovery for HCC patients. Statistical analysis and interpretable machine learning techniques were combined, to comprehensively elucidate the glycosylation alteration in HCC patients. Notably, we observed an increase in core fucosylation, especially on high-branching complex glycans, consistent with previous reports(29). We also observed a decrease of galactosylation on bi-antennary glycans, with large effect size but small fold change, which is rarely reported previously. This might be a bonus of the high sensitivity of GlyTrait. Finally, through re-analyzing a published dataset of total serum N-glycosylation from visceral leishmaniasis patients, we provided a full-scale depiction of sialic acid linkage alteration, acting as a powerful complement for the original research(28), shedding more light on the disease mechanism.

One limitation of GlyTrait is that, for now only N-glycomic data could be analyzed by GlyTrait. However, the software framework of GlyTrait is highly extensible and scalable. In the future, derived traits of other types of glycans, like O-glycans, human milk oligosaccharides, and even RNA glycans could be easily added to GlyTrait workflow. Another limitation of GlyTrait is its demand for structural annotation. Up till now, structural annotation from mass spectrometry based glycomics is not easy. Efforts have been made in this fields, including glycoforest(30) and recently developed CandyCrunch(31). We believe these advances will accelerate the prevalence of GlyTrait. As a backup, GlyTrait supports compositional-based analysis, with less structural resolution. Finally, the utility of GlyTrait has only been demonstrated on MALDI data in this study, maybe a futural comprehensive benchmark on various other platforms like UPLC-FLR will further verify its scalability.

Efforts have been made to provide tailored approaches considering the unique characteristics of glycomic data. As two pioneers in this field, GlyCompare(9) and glycowork(8) both provides exquisite methods, shifting the analyzed objects from glycan abundances to motifs or substructures. GlyTrait appears as a powerful complement to these tools. While substructure-based analysis focus on subunit of glycans that might be biologically functional (e.g. Lewis structures, blood antigens, etc.), GlyTrait provides a bird’s eye view on certain structural properties (e.g. sialylation, fucosylation, branching, etc.). Furthermore, GlyTrait covers interactions among different properties. As we have shown by the VL dataset, core fucosylation degree differed within glycans with different numbers of antennas. This information could not be easily captured by substructure-based analysis. Finally, benefiting from advances in machine learning interpretation methods, GlyTrait has the capability to delve into the intricacies of glycomic data. We believe that GlyTrait, along with other glyco-informatic tools, will substantially boost the glycomics field.

## Methods

### The EPO N-glycosylation dataset collection and curation

Previously published data(17) for EPO N-glycoprofiles was downloaded from the journal website (Supplementary Data 1 – Input 1). Structures were acquired from the spectra annotations in the supporting information of the article and converted to condensed glycoCT format via GlycoWorkBench (version 2.1 stable, build 157) (Supplementary Data 1 – Input 2). Only glycans with structure annotations were included for GlyTrait to analyze. Missing values were imputed by zero and total abundance normalization was performed in GlyTrait.

### The VL N-glycosylation dataset collection and curation

Previously published data(28) for total serum N-glycoprofiles in visceral leishmaniasis patients and controls were used to demonstrate the sialic acid linkage functionalities in this article. The final curated and normalized data (Table S5 of the original paper) was downloaded from the journal website. Only the discovery cohort was used (“SampleSet” is “Discovery”). Followed by the advice of the original paper, we excluded the non-endemic controls to exclude bias introduced by demographic difference (“Group_letter” is “Visceral Leishmaniasis”, “Asymptomatic”, or “Healthy from Endemic area”). Glycan structures were assigned according to a serum N-glycan structure library reported previously(25). For rare cases when one composition had more than one possible structure assignments, the one with larger average abundance in human serum was adopted. The data were then curated into the format required by GlyTrait. As the data itself is well-preprocessed with no missing values and with proper normalization, derived traits could be calculated directly by GlyTrait. The ANOVA results provided by GlyTrait is not used directly in this study. Instead, we performed two Welch *t*-tests (R rstatix package, v0.7.2) between VL and Asy, and between VL and EC, followed by Benjamini-Hochberg correction, for the difference between Asy and EC is irrelevant to the scientific problem here.

### The simulated serum dataset

100 simulated samples (the baseline samples) were generated for 76 serum glycans with structure annotations reported previously(25). Glycan abundances were randomly generated with the following constrains: 1) the average abundances M= {*µ*_l_, *µ*_2_,…, *µ_i_*, …, *µ*_76_} of all glycans obey a log-normal distribution with *µ* = 0 and *σ* = 1; 2) the abundances of glycan *i* obey a log-normal distribution with *µ* = *µ_i_* and *σ* = 1. To simulate core fucosylation regulation, the abundances of all core-fucosylated glycans were multiplied by a factor of 1.25, 1.50, 1.75 and 2.00 respectively. Welch *t*-test with BH adjustment were performed between the baseline samples and each regulation group, on glycan abundances and derived trait values respectively.

### Serum samples for HCC patients and controls

Serum samples were acquired from Guangdong Provincial People’s Hospital during April 2023 to June 2023. The use of human serum samples was approved by the Laboratory Medicine of Guangdong Provincial People’s Hospital and Guangdong Academy of Medical Sciences. Entry criteria of HCC patients included: (1) being diagnosed as HCC by histology; (2) no other associated malignancies; (3) absence of macroscopic hepatic vein tumor thrombus, extrahepatic spread, or distant metastases. Control samples included serum from patients with chronic hepatitis B and healthy individuals in routine medical examinations. Serum samples were stored at −80 _ before usage.

### Sample pre-processing and mass spectrometry analysis

Sample preprocessing was performed according to the methods described previously(32). Briefly, 1 μL of serum was denatured at 100 °C. N-glycans were then released by PNGase F at 37 °C overnight. Released glycans were enriched by cotton before being lyophilized. Methylamidation was then performed to protect the sialic acids. The derived glycans were enriched by cotton again and lyophilized. Samples were analyzed in a random order to mitigate batch effect. Glycan samples were stored at −20 °C before analyzed by mass spectrometry.

Sample aliquots were carefully spotted onto a MALDI target plate and left to dry at room temperature. Subsequently, 1 *µ*L of the 2,5-dihydroxybenzoic acid (DHB) matrix solution (5 mg/mL in 50% acetonitrile (ACN) with 0.1% trifluoroacetic acid (TFA)) was applied to the dried sample spots and allowed to crystallize prior to analysis. MALDI-MS spectra were recorded in the positive ion reflector mode, scanning a mass range from m/z 1000 to 4000, utilizing a rapifleX MALDI TOF mass spectrometer (Bruker). The instrument settings and calibration were optimized for the highest accuracy and resolution of glycan molecular ions.

### Data analysis for serum N-glycome data

The MALDI-TOF spectra were first processed in flexAnalysis, where the mass lists were exported to XLSX files after baseline subtraction. The annotation of the mass list files was performed using an in-house tool GlyHunter (https://github.com/FudanLuLab/glyhunter) with the retrosynthetic glycan library developed previously(33). Structures were assigned according to a serum N-glycan structure library reported previously(25) (Supplementary Data – Input 2). Compositions without structure assignment were ruled out.

The derived traits were calculated by GlyTrait with the following settings: missing values imputed by the minimum value of a glycan across samples, glycans with missing values in more than 50% samples removed, low-variance filtering on, collinearity filtering setting to 0.9.

### Pre-processing of glycoprofiles data

Three procedures were used for pre-processing the glycoprofiles before derived traits calculation. First, glycans with missing value proportions larger than certain threshold are ruled out. Second, missing values were imputed using one of the following values: mean, median, minimum or detection limit of each glycans across samples, or zero. Detection limit is defined as 0.2 × the minimum abundance of the glycan across samples. Finally, total abundance normalization (TAN) is performed on the imputed abundance table.

### The derived trait calculation and post-filtering algorithm of GlyTrait

The calculation of derived traits consists of three steps. First, the glycan abundance table is preprocessed as above, and all provided files (structural or compositional strings, built-in or custom formulas) are parsed. Then, if a custom meta property table is not provided, GlyTrait extracts N-glycan meta-properties from the structural or compositions strings. Finally, derived traits are calculated according to the formulas, along with the meta-properties extracted. The computation of derived traits leverages numpy (v1.25.2) and pandas (v2.0.1) package for vectorization. For a detailed description about derived trait calculation, please refer to Supplementary Note 1 and 2.

After derived trait calculation, GlyTrait continues with a post-filtering step. First, all derived traits with zero variance are removed. Then, the collinearity filtering algorithm removes traits with information redundancy. Two networks are built to perform the collinearity filtering: one ontology network based on trait definitions and one correlation network based on the actual correlation relationships among traits. The two networks are merged with all traits as nodes and common edges in the two networks kept. The resulting merged network is a directed acyclic graph (DAG). For each component in the DAG, only traits with an in-degree of zero are kept. For a detailed description about building the two networks and network merging, please refer to Supplementary Note 3 and 4.

### Statistical analysis

GlyTrait uses the pingouin package (v0.5.3) for statistical analysis, which is a thin wrapper around scipy.stats. For dataset with two groups, Welch *t*-test will be used. Cohen’s *d* will also be calculated as effect sizes. For dataset with more than two groups, one-way Welch ANOVA and Games-Howell post-hoc test will be used, with partial *η*^2^ as effect sizes. All *p*-values are adjusted by Benjamini-Hochberg (FDR) method. Derived traits with FDR < 0.05 were regarded significant in this study.

Statistical analysis outside GlyTrait package for data analysis in this paper was carried out using R rstatix package (v0.7.2). ROC analysis for selected derived traits were performed with R pROC package (v1.18.5) and rsample package (v1.2.0). The ROC AUC values were estimated by performing 1000 bootstrap samples, and the corresponding confidence intervals were calculated based on these sampling results. The ROC curves were obtained by overlapping the results of 100 bootstrap samples.

### Interpretable machine learning strategy

Machine learning was performed by Python scikit-learn library (v1.3.2). A nested cross-validation strategy is used for robust model evaluation and Shapley value calculation (Supplementary Fig. 11). The outer layer is 5-fold stratified repeated cross-validation, repeated for 10 times. The inner layer is 3-fold stratified cross-validation. The samples are randomly shuffled before each train-test split. In the inner layer, Python optuna library (v3.5.0) is used to fine-tune the XGBoost classifier. The best hyper-parameter set is obtained by Tree-structured Parzen Estimator algorithm (TPE) in 50 trials. Each hyper-parameter set is evaluation by 3-fold stratified cross-validation (the inner layer) using average ROC AUC across splits. In each split of the outer layer, a model is first fine-tuned in the procedure above, then trained on whole train data. The train model is then evaluated on the test fold using ROC AUC.

The Shapley values for samples in the test fold are then calculated using the “TreeExplainer” of Python SHAP library (v0.44.0). Shapley values were used to explain the probability output of the classification models in this study, instead of the default Log Odds output. The repeated cross-validation strategy (10 repeat times) of the outer layer produced 10 probabilities for each sample and 10 Shapley values for each trait in each sample. The final probability is calculated by averaging all probabilities of a sample, and the final Shapley value of a trait in a sample is calculated by averaging all Shapley values of the trait in the sample. The feature importance of a derived trait is calculated by averaging all absolute Shapley values of the trait across samples. Note that the importance of a feature group is not determined by summing the average absolute Shapley values of the features. Instead, Shapley values of each feature group in each sample are calculated first, and then the importance of the feature group according to the method applied to individual features is determined.

To assess whether the feature importance obtained from SHAP is due to random effects, we evaluate its significance through a variant of permutation tests. The labels of the data are shuffled to generate null data, which eliminates all relationships between features and labels. Null models are trained on this null data to obtain null Shapley values. Specifically, the method for calculating Shapley values is similar to the above (5-fold cross-validation, 10 repetitions, averaging the Shapley values for each sample). The method for calculating feature importance is the same as described earlier. This permutation is repeated 1000 times, yielding a distribution of null importance for each feature. The null hypothesis posits that the contribution of a feature is not attributable to random effect. In other words, the feature’s contribution in the actual model does not exceed what would be expected in a model trained on random data. The significance of this contribution, denoted by *p*_*perm*_, is calculated as the proportion of the null importance distribution that exceeds the observed importance of the feature. Finally, the *p*-values of all features are corrected through the BH method, with *p*_*perm*_ < 0.05 considered as significant. Note that since the smallest meaningful p-value we could get is 1/1000 = 0.001, when the actual calculated *p*-value is 0 (when the actual importance greater than all 1000 permutations’ importance), it is reported in the text as *p*_*perm*_ < 0.001.

### Data visualization

Results were visualized mostly using R (v4.3.2) ggplot2 package (v3.4.4) in tidyverse (v2.0.0). The sunburst plot (Fig. 1c) was created by R Plotly package (v4.10.4). The correlograms (Fig. 2g, Supplementary Fig. 2) were created by R corrplot package (v0.92). The collinearity filtering network (Fig. 3d) was created by R ggraph package (v2.1.0). The ontology networks for built-in derived traits (Supplementary Fig. 3, 4) were created with Cytoscape (v3.10.1). The heatmap for derived traits in the HCC dataset (Fig. 4a) and the VL dataset (Fig. 6a) was created with R ComplexHeatmap package (v2.18.0). The radar plots (Fig. 4b, c) were created with R ggradar package (v0.2).

## Data availability

The data generated in this study are included in this published article and its supplementary information files. The previously published glycoengineering dataset is available through https://www.nature.com/articles/nbt.3280#Sec6. The previously published visceral leishmaniasis dataset is available through glycoPOST with accession ID: GPST000313 [https://glycopost.glycosmos.org/entry/GPST000313]. The raw data of the hepatocellular carcinoma dataset is available upon request. Source data are provided with this paper.

## Code availability

GlyTrait was developed in Python 3.10. The source code of GlyTrait could be accessed from the GitHub page under an MIT license: https://github.com/FudanLuLab/glytrait. The command line interface (CLI) and the Python package could be installed from PyPi (https://pypi.org/project/glytrait). In addition to the command line tool, a web-application built with the Streamlit library (v1.25.0) is available through https://glytrait.streamlit.app. To keep GlyTrait lightweight, currently the ML workflow is provided as Jupyter Notebooks, available in the same depository as the CLI.

## Acknowledgements

This work is supported by the National Key Research and Development Program of China (2021YFA1301602), the National Natural Science Foundation of China (Grants 22174021 and 82121004), Shanghai Project (22142202400).

