## Supplementary Information for "GlyTrait Brings Insights into Functional Glycosylation"

### GlyTrait: A versatile bioinformatics tool for Glycomics Analysis

### Supplementary Tables

### Supplementary Notes

### Supplementary Figures

### Supplementary Table 1 – Built-in structural derived traits

In the table, “x” can be 1, 2, 3, or 4, meaning mono-, bi-, tri-, or tetra-antennary glycans, respectively.

For example, “CAx” can be “CA1”, “CA2”, “CA3”, and “CA4”, meaning “the relative abundance of mono-, bi-, tri-, or tetra-antennary glycans within complex-type glycans”, and the formula could be materialized into “CA1 = [nAnt == 1] // [type == ‘complex’]”, etc.

| **Trait name** | **Definition** |
| --- | --- |
|  | **Formula expression** |
| TM | Relative abundance of high-mannose type glycans within total spectrum |
|  | TM = [type == 'high-mannose'] / [1] |
| THy | Relative abundance of hybrid type glycans within total spectrum |
|  | THy = [type == 'hybrid'] / [1] |
| MHy | The ratio of high-mannose to hybrid glycans |
|  | MHy = [type == 'high-mannose'] / [type == 'hybrid'] |
| MM | Average number of mannoses on high mannose type glycans |
|  | MM = [nM] // [type == 'high-mannose'] |
| CAx | Relative abundance of x-antennary glycans within complex-type glycans |
|  | CAx = [nAnt == x] // [type == 'complex'] |
| CFc | Core fucosylation within complex type glycans |
|  | CFc = [nFc > 0] // [type == 'complex'] |
| AxFc | Core fucosylation within x-antennary glycans |
|  | AxFc = [nFc > 0] // [(type == 'complex') * (nAnt == x)] |
| AxSFc | Core fucosylation within sialylated x-antennary glycans |
|  | AxSFc = [nFc > 0] // [(type == 'complex') * (nAnt == x) * (nS > 0)] |
| AxS0Fc | Core fucosylation within asialylated x-antennary glycans |
|  | AxS0Fc = [nFc > 0] // [(type == 'complex') * (nAnt == x) * (nS == 0)] |
| AxEFc | Core fucosylation within x-antennary glycans with α2,6-linked sialic acids |
|  | AxEFc = [nFc > 0] // [(type == 'complex') * (nAnt == x) * (nE > 0)] |
| AxE0Fc | Core fucosylation within x-antennary glycans without α2,6-linked sialic acids |
|  | AxE0Fc = [nFc > 0] // [(type == 'complex') * (nAnt == x) * (nE == 0)] |
| AxLFc | Core fucosylation within x-antennary glycans with α2,3-linked sialic acids |
|  | AxLFc = [nFc > 0] // [(type == 'complex') * (nAnt == x) * (nL > 0)] |
| AxL0Fc | Core fucosylation within x-antennary glycans without α2,3-linked sialic acids |
|  | AxL0Fc = [nFc > 0] // [(type == 'complex') * (nAnt == x) * (nL == 0)] |
| CFa | Antennary fucosylation within complex type glycans |
|  | CFa = [nFa] // [type == 'complex'] |
| AxFa | Antennary fucosylation within x-antennary glycans |
|  | AxFa = [nFa] // [(type == 'complex') * (nAnt == x)] |
| AxSFa | Antennary fucosylation within sialylated x-antennary glycans |
|  | AxSFa = [nFa] // [(type == 'complex') * (nAnt == x) * (nS > 0)] |
| AxS0Fa | Antennary fucosylation within asialylated x-antennary glycans |
|  | AxS0Fa = [nFa] // [(type == 'complex') * (nAnt == x) * (nS == 0)] |
| AxEFa | Antennary fucosylation within x-antennary glycans with α2,6-linked sialic acids |
|  | AxEFa = [nFa] // [(type == 'complex') * (nAnt == x) * (nE > 0)] |
| AxE0Fa | Antennary fucosylation within x-antennary glycans without α2,6-linked sialic acids |
|  | AxE0Fa = [nFa] // [(type == 'complex') * (nAnt == x) * (nE == 0)] |
| AxLFa | Antennary fucosylation within x-antennary glycans with α2,3-linked sialic acids |
|  | AxLFa = [nFa] // [(type == 'complex') * (nAnt == x) * (nL > 0)] |
| AxL0Fa | Antennary fucosylation within x-antennary glycans without α2,3-linked sialic acids |
|  | AxL0Fa = [nFa] // [(type == 'complex') * (nAnt == x) * (nL == 0)] |
| CB | Relative abundance of bisected glycans within all complex glycans |
|  | CB = [B] // [type == 'complex'] |
| AxB | Relative abundance of bisected glycans within x-antennary glycans |
|  | AxB = [B] // [(type == 'complex') * (nAnt == x)] |
| AxFB | Relative abundance of bisected glycans within fucosylated x-antennary glycans |
|  | AxFB = [B] // [(type == 'complex') * (nAnt == x) * (nF > 0)] |
| AxF0B | Relative abundance of bisected glycans within afucosylated x-antennary glycans |
|  | AxF0B = [B] // [(type == 'complex') * (nAnt == x) * (nF == 0)] |
| AxSB | Relative abundance of bisected glycans within sialylated x-antennary glycans |
|  | AxSB = [B] // [(type == 'complex') * (nAnt == x) * (nS > 0)] |
| AxS0B | Relative abundance of bisected glycans within asialylated x-antennary glycans |
|  | AxS0B = [B] // [(type == 'complex') * (nAnt == x) * (nS == 0)] |
| AxFSB | Relative abundance of bisected glycans within sialylated and fucosylated x-antennary glycans |
|  | AxFSB = [B] // [(type == 'complex') * (nAnt == x) * (nF > 0) * (nS > 0)] |
| AxF0SB | Relative abundance of bisected glycans within sialylated and afucosylated x-antennary glycans |
|  | AxF0SB = [B] // [(type == 'complex') * (nAnt == x) * (nF == 0) * (nS > 0)] |
| AxFS0B | Relative abundance of bisected glycans within asialylated and fucosylated x-antennary glycans |
|  | AxFS0B = [B] // [(type == 'complex') * (nAnt == x) * (nF > 0) * (nS == 0)] |
| AxF0S0B | Relative abundance of bisected glycans within asialylated and afucosylated x-antennary glycans |
|  | AxF0S0B = [B] // [(type == 'complex') * (nAnt == x) * (nF == 0) * (nS == 0)] |
| AxLB | Relative abundance of bisected glycans within x-antennary glycans with α2,3-linked sialic acids |
|  | AxLB = [B] // [(type == 'complex') * (nAnt == x) * (nL > 0)] |
| AxL0B | Relative abundance of bisected glycans within x-antennary glycans without α2,3-linked sialic acids |
|  | AxL0B = [B] // [(type == 'complex') * (nAnt == x) * (nL == 0)] |
| AxFLB | Relative abundance of bisected glycans within fucosylated x-antennary glycans with α2,3-linked sialic acids |
|  | AxFLB = [B] // [(type == 'complex') * (nAnt == x) * (nL > 0) * (nF > 0)] |
| AxF0LB | Relative abundance of bisected glycans within afucosylated x-antennary glycans with α2,3-linked sialic acids |
|  | AxF0LB = [B] // [(type == 'complex') * (nAnt == x) * (nL > 0) * (nF == 0)] |
| AxFL0B | Relative abundance of bisected glycans within fucosylated x-antennary glycans without α2,3-linked sialic acids |
|  | AxFL0B = [B] // [(type == 'complex') * (nAnt == x) * (nF > 0) * (nL == 0)] |
| AxF0L0B | Relative abundance of bisected glycans within afucosylated x-antennary glycans without α2,3-linked sialic acids |
|  | AxF0L0B = [B] // [(type == 'complex') * (nAnt == x) * (nF == 0) * (nL == 0)] |
| AxEB | Relative abundance of bisected glycans within x-antennary glycans with α2,6-linked sialic acids |
|  | AxEB = [B] // [(type == 'complex') * (nAnt == x) * (nE > 0)] |
| AxE0B | Relative abundance of bisected glycans within x-antennary glycans without α2,6-linked sialic acids |
|  | AxE0B = [B] // [(type == 'complex') * (nAnt == x) * (nE > 0)] |
| AxFEB | Relative abundance of bisected glycans within fucosylated x-antennary glycans with α2,6-linked sialic acids |
|  | AxFEB = [B] // [(type == 'complex') * (nAnt == x) * (nF > 0) * (nE > 0)] |
| AxF0EB | Relative abundance of bisected glycans within afucosylated x-antennary glycans with α2,6-linked sialic acids |
|  | AxF0EB = [B] // [(type == 'complex') * (nAnt == x) * (nF == 0) * (nE > 0)] |
| AxFE0B | Relative abundance of bisected glycans within fucosylated x-antennary glycans without α2,6-linked sialic acids |
|  | AxFE0B = [B] // [(type == 'complex') * (nAnt == x) * (nF > 0) * (nE == 0)] |
| AxF0E0B | Relative abundance of bisected glycans within afucosylated x-antennary glycans without α2,6-linked sialic acids |
|  | AxF0E0B = [B] // [(type == 'complex') * (nAnt == x) * (nF == 0) * (nE == 0)] |
| CG | Average number of galactoses on complex glycans |
|  | CG = [nG] // [type == 'complex'] |
| AxG | Galactosylation per antenna within x-antennary glycans |
|  | AxG = [nG / x] // [(type == 'complex') * (nAnt == x)] |
| AxFG | Galactosylation per antenna within fucosylated x-antennary glycans |
|  | AxFG = [nG / x] // [(type == 'complex') * (nAnt == x) * (nF > 0)] |
| AxF0G | Galactosylation per antenna within afucosylated x-antennary glycans |
|  | AxF0G = [nG / x] // [(type == 'complex') * (nAnt == x) * (nF == 0)] |
| AxSG | Galactosylation per antenna within sialylated x-antennary glycans |
|  | AxSG = [nG / x] // [(type == 'complex') * (nAnt == x) * (nS > 0)] |
| AxS0G | Galactosylation per antenna within asialylated x-antennary glycans |
|  | AxS0G = [nG / x] // [(type == 'complex') * (nAnt == x) * (nS == 0)] |
| AxFSG | Galactosylation per antenna within fucosylated sialylated x-antennary glycans |
|  | AxFSG = [nG / x] // [(type == 'complex') * (nAnt == x) * (nF > 0) * (nS > 0)] |
| AxF0SG | Galactosylation per antenna within afucosylated sialylated x-antennary glycans |
|  | AxF0SG = [nG / x] // [(type == 'complex') * (nAnt == x) * (nF == 0) * (nS > 0)] |
| AxFS0G | Galactosylation per antenna within fucosylated asialylated x-antennary glycans |
|  | AxFS0G = [nG / x] // [(type == 'complex') * (nAnt == x) * (nF > 0) * (nS == 0)] |
| AxF0S0G | Galactosylation per antenna within afucosylated asialylated x-antennary glycans |
|  | AxF0S0G = [nG / x] // [(type == 'complex') * (nAnt == x) * (nF == 0) * (nS == 0)] |
| AxLG | Galactosylation per antenna within α2,3-sialylated x-antennary glycans |
|  | AxLG = [nG / x] // [(type == 'complex') * (nAnt == x) * (nL > 0)] |
| AxL0G | Galactosylation per antenna within x-antennary glycans without α2,3-linked sialic acids |
|  | AxL0G = [nG / x] // [(type == 'complex') * (nAnt == x) * (nL == 0)] |
| AxFLG | Galactosylation per antenna within fucosylated α2,3-sialylated x-antennary glycans |
|  | AxFLG = [nG / x] // [(type == 'complex') * (nAnt == x) * (nF > 0) * (nL > 0)] |
| AxF0LG | Galactosylation per antenna within afucosylated α2,3-sialylated x-antennary glycans |
|  | AxF0LG = [nG / x] // [(type == 'complex') * (nAnt == x) * (nF == 0) * (nL > 0)] |
| AxFL0G | Galactosylation per antenna within fucosylated x-antennary glycans without α2,3-linked sialic acids |
|  | AxFL0G = [nG / x] // [(type == 'complex') * (nAnt == x) * (nF > 0) * (nL == 0)] |
| AxF0L0G | Galactosylation per antenna within afucosylated x-antennary glycans without α2,3-linked sialic acids |
|  | AxF0L0G = [nG / x] // [(type == 'complex') * (nAnt == x) * (nF == 0) * (nL == 0)] |
| AxEG | Galactosylation per antenna within α2,6-sialylated x-antennary glycans |
|  | AxEG = [nG / x] // [(type == 'complex') * (nAnt == x) * (nE > 0)] |
| AxE0G | Galactosylation per antenna within x-antennary glycans without α2,6-linked sialic acids |
|  | AxE0G = [nG / x] // [(type == 'complex') * (nAnt == x) * (nE == 0)] |
| AxFEG | Galactosylation per antenna within fucosylated α2,6-sialylated x-antennary glycans |
|  | AxFEG = [nG / x] // [(type == 'complex') * (nAnt == x) * (nF > 0) * (nE > 0)] |
| AxF0EG | Galactosylation per antenna within afucosylated α2,6-sialylated x-antennary glycans |
|  | AxF0EG = [nG / x] // [(type == 'complex') * (nAnt == x) * (nF == 0) * (nE > 0)] |
| AxFE0G | Galactosylation per antenna within fucosylated x-antennary glycans without α2,6-linked sialic acids |
|  | AxFE0G = [nG / x] // [(type == 'complex') * (nAnt == x) * (nF > 0) * (nE == 0)] |
| AxF0E0G | Galactosylation per antenna within afucosylated x-antennary glycans without α2,6-linked sialic acids |
|  | AxF0E0G = [nG / x] // [(type == 'complex') * (nAnt == x) * (nF == 0) * (nE == 0)] |
| CS | Average number of sialic acids on complex glycans |
|  | CS = [nS] // [type == 'complex'] |
| AxS | Sialylation per antenna within x-antennary glycans |
|  | AxS = [nS / x] // [(type == 'complex') * (nAnt == x)] |
| AxFS | Sialylation per antenna within fucosylated x-antennary glycans |
|  | AxFS = [nS / x] // [(type == 'complex') * (nAnt == x) * (nF > 0)] |
| AxF0S | Sialylation per antenna within afucosylated x-antennary glycans |
|  | AxF0S = [nS / x] // [(type == 'complex') * (nAnt == x) * (nF == 0)] |
| AxGS | Sialylation per galactose within x-antennary glycans |
|  | AxGS = [nS / nG] // [(type == 'complex') * (nAnt == x)] |
| AxFGS | Sialylation per galactose within fucosylated x-antennary glycans |
|  | AxFGS = [nS / nG] // [(type == 'complex') * (nAnt == x) * (nF > 0)] |
| AxF0GS | Sialylation per galactose within afucosylated x-antennary glycans |
|  | AxF0GS = [nS / nG] // [(type == 'complex') * (nAnt == x) * (nF == 0)] |
| CL | Average number of α2,3-linked sialic acids on complex glycans |
|  | CL = [nL] // [type == 'complex'] |
| AxL | α2,3-sialylation per antenna within x-antennary glycans |
|  | AxL = [nL / x] // [(type == 'complex') * (nAnt == x)] |
| AxFL | α2,3-sialylation per antenna within fucosylated x-antennary glycans |
|  | AxFL = [nL / x] // [(type == 'complex') * (nAnt == x) * (nF > 0)] |
| AxF0L | α2,3-sialylation per antenna within afucosylated x-antennary glycans |
|  | AxF0L = [nL / x] // [(type == 'complex') * (nAnt == x) * (nF == 0)] |
| AxGL | α2,3-sialylation per galactose within x-antennary glycans |
|  | AxGL = [nL / nG] // [(type == 'complex') * (nAnt == x)] |
| AxFGL | α2,3-sialylation per galactose within fucosylated x-antennary glycans |
|  | AxFGL = [nL / nG] // [(type == 'complex') * (nAnt == x) * (nF > 0)] |
| AxF0GL | α2,3-sialylation per galactose within afucosylated x-antennary glycans |
|  | AxF0GL = [nL / nG] // [(type == 'complex') * (nAnt == x) * (nF == 0)] |
| CE | Average number of α2,6-linked sialic acids on complex glycans |
|  | CE = [nE] // [type == 'complex'] |
| AxE | α2,6-sialylation per antenna within x-antennary glycans |
|  | AxE = [nE / x] // [(type == 'complex') * (nAnt == x)] |
| AxFE | α2,6-sialylation per antenna within fucosylated x-antennary glycans |
|  | AxFE = [nE / x] // [(type == 'complex') * (nAnt == x) * (nF > 0)] |
| AxF0E | α2,6-sialylation per antenna within afucosylated x-antennary glycans |
|  | AxF0E = [nE / x] // [(type == 'complex') * (nAnt == x) * (nF == 0)] |
| AxGE | α2,6-sialylation per galactose within x-antennary glycans |
|  | AxGE = [nE / nG] // [(type == 'complex') * (nAnt == x)] |
| AxFGE | α2,6-sialylation per galactose within fucosylated x-antennary glycans |
|  | AxFGE = [nE / nG] // [(type == 'complex') * (nAnt == x) * (nF > 0)] |
| AxF0GE | α2,6-sialylation per galactose within afucosylated x-antennary glycans |
|  | AxF0GE = [nE / nG] // [(type == 'complex') * (nAnt == x) * (nF == 0)] |
| CPl | Relative abundance of glycans with poly-LacNAc within all complex glycans |
|  | CPl = [Pl] // [type == 'complex'] |
| AxPl | Relative abundance of glycans with poly-LacNAc within x-antennary glycans |
|  | AxPl = [Pl] // [(type == 'complex') * (nAnt == x)] |
| AxFPl | Relative abundance of glycans with poly-LacNAc within fucosylated x-antennary glycans |
|  | AxFPl = [Pl] // [(type == 'complex') * (nAnt == x) * (nF > 0)] |
| AxF0Pl | Relative abundance of glycans with poly-LacNAc within afucosylated x-antennary glycans |
|  | AxF0Pl = [Pl] // [(type == 'complex') * (nAnt == x) * (nF == 0)] |
| AxSPl | Relative abundance of glycans with poly-LacNAc within sialylated x-antennary glycans |
|  | AxSPl = [Pl] // [(type == 'complex') * (nAnt == x) * (nS > 0)] |
| AxS0Pl | Relative abundance of glycans with poly-LacNAc within asialylated x-antennary glycans |
|  | AxS0Pl = [Pl] // [(type == 'complex') * (nAnt == x) * (nS == 0)] |
| AxFSPl | Relative abundance of species with poly-LacNAc within fucosylated sialylated x-antennary glycans |
|  | AxFSPl = [Pl] // [(type == 'complex') * (nAnt == x) * (nF > 0) * (nS > 0)] |
| AxF0SPl | Relative abundance of species with poly-LacNAc within afucosylated sialylated x-antennary glycans |
|  | AxF0SPl = [Pl] // [(type == 'complex') * (nAnt == x) * (nF == 0) * (nS > 0)] |
| AxFS0Pl | Relative abundance of species with poly-LacNAc within fucosylated asialylated x-antennary glycans |
|  | AxF0SPl = [Pl] // [(type == 'complex') * (nAnt == x) * (nF > 0) * (nS == 0)] |
| AxF0S0Pl | Relative abundance of species with poly-LacNAc within afucosylated asialylated x-antennary glycans |
|  | AxF0S0Pl = [Pl] // [(type == 'complex') * (nAnt == x) * (nF == 0) * (nS == 0)] |
| AxEPl | Relative abundance of glycans with poly-LacNAc within α2,6-sialylated x-antennary glycans |
|  | AxEPl = [Pl] // [(type == 'complex') * (nAnt == x) * (nE > 0)] |
| AxE0Pl | Relative abundance of glycans with poly-LacNAc x-antennary glycans without α2,6-linked sialic acids |
|  | AxE0Pl = [Pl] // [(type == 'complex') * (nAnt == x) * (nE == 0)] |
| AxFEPl | Relative abundance of species with poly-LacNAc within fucosylated α2,6-sialylated x-antennary glycans |
|  | AxFEPl = [Pl] // [(type == 'complex') * (nAnt == x) * (nF > 0) * (nE > 0)] |
| AxF0EPl | Relative abundance of species with poly-LacNAc within afucosylated α2,6-sialylated x-antennary glycans |
|  | AxF0EPl = [Pl] // [(type == 'complex') * (nAnt == x) * (nF == 0) * (nE > 0)] |
| AxFE0Pl | Relative abundance of species with poly-LacNAc within fucosylated x-antennary glycans without α2,6-linked sialic acids |
|  | AxF0EPl = [Pl] // [(type == 'complex') * (nAnt == x) * (nF > 0) * (nE == 0)] |
| AxF0E0Pl | Relative abundance of species with poly-LacNAc within afucosylated x-antennary glycans without α2,6-linked sialic acids |
|  | AxF0E0Pl = [Pl] // [(type == 'complex') * (nAnt == x) * (nF == 0) * (nE == 0)] |
| AxLPl | Relative abundance of glycans with poly-LacNAc within α2,3-sialylated x-antennary glycans |
|  | AxLPl = [Pl] // [(type == 'complex') * (nAnt == x) * (nL > 0)] |
| AxL0Pl | Relative abundance of glycans with poly-LacNAc x-antennary glycans without α2,3-linked sialic acids |
|  | AxL0Pl = [Pl] // [(type == 'complex') * (nAnt == x) * (nL == 0)] |
| AxFLPl | Relative abundance of species with poly-LacNAc within fucosylated α2,3-sialylated x-antennary glycans |
|  | AxFLPl = [Pl] // [(type == 'complex') * (nAnt == x) * (nF > 0) * (nL > 0)] |
| AxF0LPl | Relative abundance of species with poly-LacNAc within afucosylated α2,3-sialylated x-antennary glycans |
|  | AxF0LPl = [Pl] // [(type == 'complex') * (nAnt == x) * (nF == 0) * (nL > 0)] |
| AxFL0Pl | Relative abundance of species with poly-LacNAc within fucosylated x-antennary glycans without α2,3-linked sialic acids |
|  | AxF0LPl = [Pl] // [(type == 'complex') * (nAnt == x) * (nF > 0) * (nL == 0)] |
| AxF0L0Pl | Relative abundance of species with poly-LacNAc within afucosylated x-antennary glycans without α2,3-linked sialic acids |
|  | AxF0L0Pl = [Pl] // [(type == 'complex') * (nAnt == x) * (nF == 0) * (nL == 0)] |

### Supplementary Table 2 – Built-in compositional derived traits

| **Trait name** | **Definition** |
| --- | --- |
|  | **Formula expression** |
| Hb | Relative abundance of high-branching (N > 4) glycans within all glycans |
|  | Hb = [nN > 4] / [1] |
| Lb | Relative abundance of low-branching (N <= 4) glycans within all glycans |
|  | Lb = [nN <= 4] / [1] |
| F | Fucosylation within all glycans |
|  | F = [nF > 0] / [1] |
| HbF | Fucosylation within high-branching glycans |
|  | HbF = [nF > 0] // [nN > 4] |
| LbF | Fucosylation within low-branching glycans |
|  | HbF = [nF > 0] // [nN <= 4] |
| HbSF | Fucosylation within sialylated high-branching glycans |
|  | HbSF = [nF > 0] // [(nN > 4) * (nS > 0)] |
| LbSF | Fucosylation within sialylated low-branching glycans |
|  | LbSF = [nF > 0] // [(nN <= 4) * (nS > 0)] |
| HbS0F | Fucosylation within asialylated high-branching glycans |
|  | HbS0F = [nF > 0] // [(nN > 4) * (nS == 0)] |
| LbS0F | Fucosylation within asialylated low-branching glycans |
|  | LbS0F = [nF > 0] // [(nN <= 4) * (nS == 0)] |
| HbEF | Fucosylation within high-branching glycans with a2,6-linked sialic acids |
|  | HbEF = [nF > 0] // [(nN > 4) * (nE > 0)] |
| LbEF | Fucosylation within low-branching glycans with a2,6-linked sialic acids |
|  | LbEF = [nF > 0] // [(nN <= 4) * (nE > 0)] |
| HbE0F | Fucosylation within high-branching glycans without a2,6-linked sialic acids |
|  | HbE0F = [nF > 0] // [(nN > 4) * (nE == 0)] |
| LbE0F | Fucosylation within low-branching glycans without a2,6-linked sialic acids |
|  | LbE0F = [nF > 0] // [(nN <= 4) * (nE == 0)] |
| HbLF | Fucosylation within high-branching glycans with a2,3-linked sialic acids |
|  | HbLF = [nF > 0] // [(nN > 4) * (nL > 0)] |
| LbLF | Fucosylation within low-branching glycans with a2,3-linked sialic acids |
|  | LbLF = [nF > 0] // [(nN <= 4) * (nL > 0)] |
| HbL0F | Fucosylation within high-branching glycans without a2,3-linked sialic acids |
|  | HbL0F = [nF > 0] // [(nN > 4) * (nL == 0)] |
| LbL0F | Fucosylation within low-branching glycans without a2,3-linked sialic acids |
|  | LbL0F = [nF > 0] // [(nN <= 4) * (nL == 0)] |
| G | Galactosylation within all glycans |
|  | G = [nG] / [1] |
| HbG | Galactosylation within high-branching glycans |
|  | HbG = [nG] // [nN > 4] |
| LbG | Galactosylation within low-branching glycans |
|  | LbG = [nG] // [nN <= 4] |
| HbFG | Galactosylation within fucosylated high-branching glycans |
|  | HbFG = [nG] // [(nN > 4) * (nF > 0)] |
| LbFG | Galactosylation within fucosylated low-branching glycans |
|  | LbFG = [nG] // [(nN <= 4) * (nF > 0)] |
| HbF0G | Galactosylation within afucosylated high-branching glycans |
|  | HbF0G = [nG] // [(nN > 4) * (nF == 0)] |
| LbF0G | Galactosylation within afucosylated low-branching glycans |
|  | LbF0G = [nG] // [(nN <= 4) * (nF == 0)] |
| HbSG | Galactosylation within sialylated high-branching glycans |
|  | HbSG = [nG] // [(nN > 4) * (nS > 0)] |
| LbSG | Galactosylation within sialylated low-branching glycans |
|  | LbSG = [nG] // [(nN <= 4) * (nS > 0)] |
| HbS0G | Galactosylation within asialylated high-branching glycans |
|  | HbS0G = [nG] // [(nN > 4) * (nS == 0)] |
| LbS0G | Galactosylation within asialylated low-branching glycans |
|  | LbS0G = [nG] // [(nN <= 4) * (nS == 0)] |
| HbFSG | Galactosylation within fucosylated sialylated high-branching glycans |
|  | HbFSG = [nG] // [(nN > 4) * (nF > 0) * (nS > 0)] |
| LbFSG | Galactosylation within fucosylated sialylated low-branching glycans |
|  | LbFSG = [nG] // [(nN <= 4) * (nF > 0) * (nS > 0)] |
| HbF0SG | Galactosylation within afucosylated sialylated high-branching glycans |
|  | HbF0SG = [nG] // [(nN > 4) * (nF == 0) * (nS > 0)] |
| LbF0SG | Galactosylation within afucosylated sialylated low-branching glycans |
|  | LbF0SG = [nG] // [(nN <= 4) * (nF == 0) * (nS > 0)] |
| HbFS0G | Galactosylation within fucosylated asialylated high-branching glycans |
|  | HbFS0G = [nG] // [(nN > 4) * (nF > 0) * (nS == 0)] |
| LbFS0G | Galactosylation within fucosylated asialylated low-branching glycans |
|  | LbFS0G = [nG] // [(nN <= 4) * (nF > 0) * (nS == 0)] |
| HbF0S0G | Galactosylation within afucosylated asialylated high-branching glycans |
|  | HbF0S0G = [nG] // [(nN > 4) * (nF == 0) * (nS == 0)] |
| LbF0S0G | Galactosylation within afucosylated asialylated low-branching glycans |
|  | LbF0S0G = [nG] // [(nN <= 4) * (nF == 0) * (nS == 0)] |
| S | Sialylation within all glycans |
|  | S = [nS] / [1] |
| HbS | Sialylation within high-branching glycans |
|  | HbS = [nS] // [nN > 4] |
| LbS | Sialylation within low-branching glycans |
|  | LbS = [nS] // [nN <= 4] |
| HbFS | Sialylation within fucosylated high-branching glycans |
|  | HbFS = [nS] // [(nN > 4) * (nF > 0)] |
| LbFS | Sialylation within fucosylated low-branching glycans |
|  | LbFS = [nS] // [(nN <= 4) * (nF > 0)] |
| HbF0S | Sialylation within afucosylated high-branching glycans |
|  | HbF0S = [nS] // [(nN > 4) * (nF == 0)] |
| LbF0S | Sialylation within afucosylated low-branching glycans |
|  | LbF0S = [nS] // [(nN <= 4) * (nF == 0)] |
| HbGS | Sialylation per galactose within high-branching glycans |
|  | HbGS = [nS / nG] // [nN > 4] |
| LbGS | Sialylation per galactose within low-branching glycans |
|  | LbGS = [nS / nG] // [nN <= 4] |
| HbFGS | Sialylation per galactose within fucosylated high-branching glycans |
|  | HbFGS = [nS / nG] // [(nN > 4) * (nF > 0)] |
| LbFGS | Sialylation per galactose within fucosylated low-branching glycans |
|  | LbFGS = [nS / nG] // [(nN <= 4) * (nF > 0)] |
| HbF0GS | Sialylation per galactose within afucosylated high-branching glycans |
|  | HbF0GS = [nS / nG] // [(nN > 4) * (nF == 0)] |
| LbF0GS | Sialylation per galactose within afucosylated low-branching glycans |
|  | LbF0GS = [nS / nG] // [(nN <= 4) * (nF == 0)] |
| L | α2,3-linked sialylation within all glycans |
|  | L = [nL] / [1] |
| HbL | α2,3-linked sialylation within high-branching glycans |
|  | HbL = [nL] // [nN > 4] |
| LbL | α2,3-linked sialylation within low-branching glycans |
|  | LbL = [nL] // [nN <= 4] |
| HbFL | α2,3-linked sialylation within fucosylated high-branching glycans |
|  | HbFL = [nL] // [(nN > 4) * (nF > 0)] |
| LbFL | α2,3-linked sialylation within fucosylated low-branching glycans |
|  | LbFL = [nL] // [(nN <= 4) * (nF > 0)] |
| HbF0L | α2,3-linked sialylation within afucosylated high-branching glycans |
|  | HbF0L = [nL] // [(nN > 4) * (nF == 0)] |
| LbF0L | α2,3-linked sialylation within afucosylated low-branching glycans |
|  | LbF0L = [nL] // [(nN <= 4) * (nF == 0)] |
| HbGL | α2,3-linked sialylation per galactose within high-branching glycans |
|  | HbGL = [nL / nG] // [nN > 4] |
| LbGL | α2,3-linked sialylation per galactose within low-branching glycans |
|  | LbGL = [nL / nG] // [nN <= 4] |
| HbFGL | α2,3-linked sialylation per galactose within fucosylated high-branching glycans |
|  | HbFGL = [nL / nG] // [(nN > 4) * (nF > 0)] |
| LbFGL | α2,3-linked sialylation per galactose within fucosylated low-branching glycans |
|  | LbFGL = [nL / nG] // [(nN <= 4) * (nF > 0)] |
| HbF0GL | α2,3-linked sialylation per galactose within afucosylated high-branching glycans |
|  | HbF0GL = [nL / nG] // [(nN > 4) * (nF == 0)] |
| LbF0GL | α2,3-linked sialylation per galactose within afucosylated low-branching glycans |
|  | LbF0GL = [nL / nG] // [(nN <= 4) * (nF == 0)] |
| E | α2,6-linked sialylation within all glycans |
|  | E = [nE] / [1] |
| HbE | α2,6-linked sialylation within high-branching glycans |
|  | HbE = [nE] // [nN > 4] |
| LbE | α2,6-linked sialylation within low-branching glycans |
|  | LbE = [nE] // [nN <= 4] |
| HbFE | α2,6-linked sialylation within fucosylated high-branching glycans |
|  | HbFE = [nE] // [(nN > 4) * (nF > 0)] |
| LbFE | α2,6-linked sialylation within fucosylated low-branching glycans |
|  | LbFE = [nE] // [(nN <= 4) * (nF > 0)] |
| HbF0E | α2,6-linked sialylation within afucosylated high-branching glycans |
|  | HbF0E = [nE] // [(nN > 4) * (nF == 0)] |
| LbF0E | α2,6-linked sialylation within afucosylated low-branching glycans |
|  | LbF0E = [nE] // [(nN <= 4) * (nF == 0)] |
| HbGE | α2,6-linked sialylation per galactose within high-branching glycans |
|  | HbGE = [nE / nG] // [nN > 4] |
| LbGE | α2,6-linked sialylation per galactose within low-branching glycans |
|  | LbGE = [nE / nG] // [nN <= 4] |
| HbFGE | α2,6-linked sialylation per galactose within fucosylated high-branching glycans |
|  | HbFGE = [nE / nG] // [(nN > 4) * (nF > 0)] |
| LbFGE | α2,6-linked sialylation per galactose within fucosylated low-branching glycans |
|  | LbFGE = [nE / nG] // [(nN <= 4) * (nF > 0)] |
| HbF0GE | α2,6-linked sialylation per galactose within afucosylated high-branching glycans |
|  | HbF0GE = [nE / nG] // [(nN > 4) * (nF == 0)] |
| LbF0GE | α2,6-linked sialylation per galactose within afucosylated low-branching glycans |
|  | LbF0GE = [nE / nG] // [(nN <= 4) * (nF == 0)] |

### Supplementary Table 3 – Glycans identified in the HCC dataset

Structures were annotated according to previously reported human serum N-glycans^1^.

| **Composition** | **Structure** | **Composition** | **Structure** |
| --- | --- | --- | --- |
| H5N4S2 | 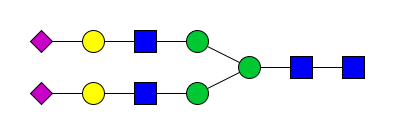 | H5N4S1 | 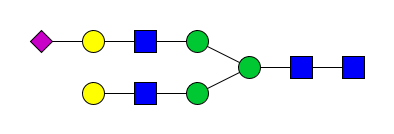 |
| H5N5F1S1 | 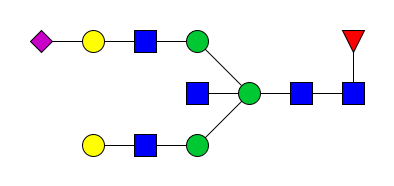 | H5N4F1 | 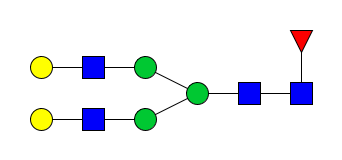 |
| H5N4 | 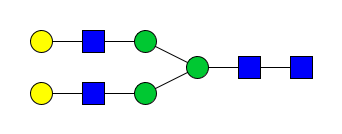 | H3N4F1 | 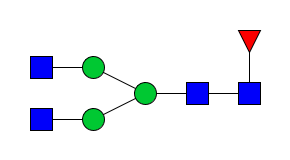 |
| H4N4F1 | 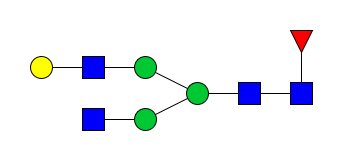 | H5N4F1S1 | 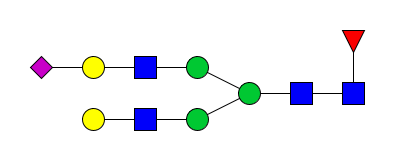 |
| H5N4F1S2 | 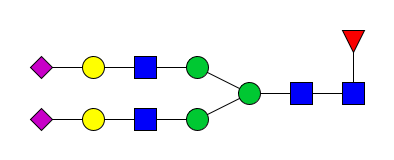 | H6N5S2 | 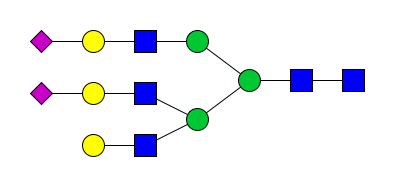 |
| H5N5F1 | 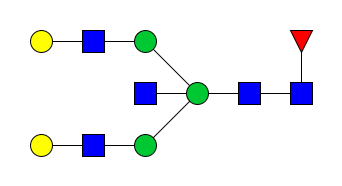 | H5N5F1S2 | 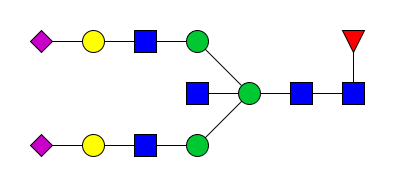 |
| H4N5F1 | 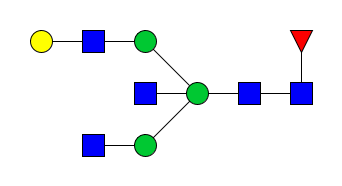 | H6N5S1 | 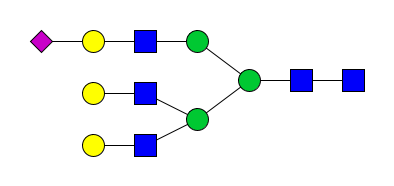 |
| H3N5F1 | 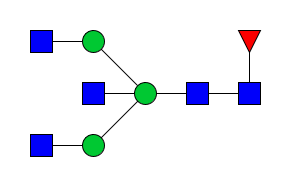 | H5N2 | 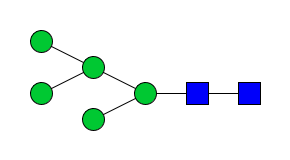 |
| H6N2 | 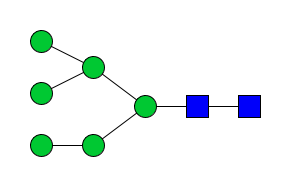 | H5N5 | 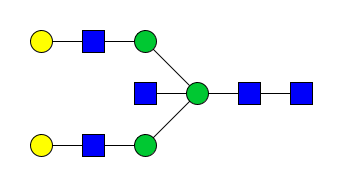 |
| H8N2 | 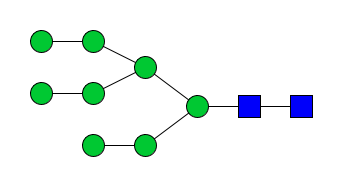 | H4N3F1 | 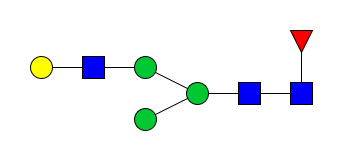 |
| H6N5F1S1 | 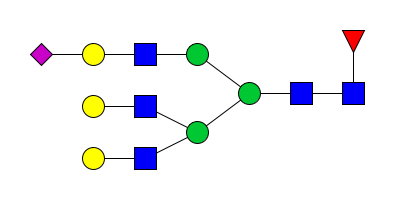 | H4N4 | 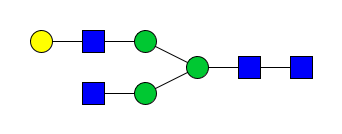 |
| H9N2 | 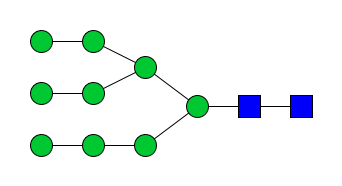 | H5N5S1 | 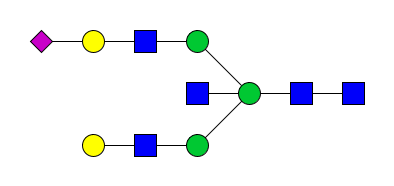 |
| H4N4S1 | 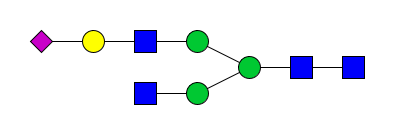 | H6N5 | 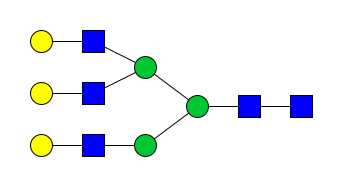 |
| H6N5F1S2 | 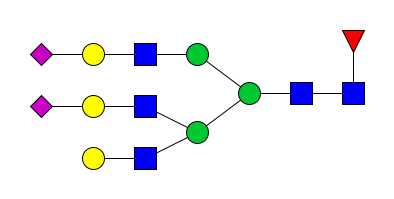 | H4N5 | 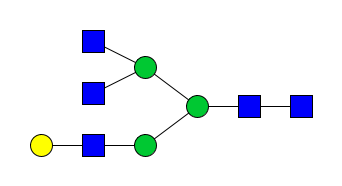 |
| H4N4F1S1 | 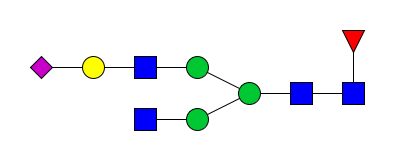 | H3N5 | 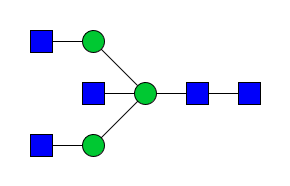 |
| H4N3S1 | 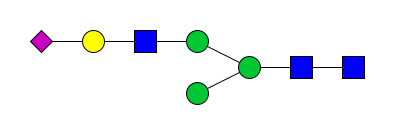 | H3N4 | 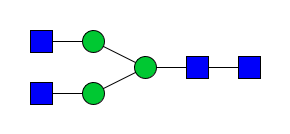 |
| H6N3S1 | 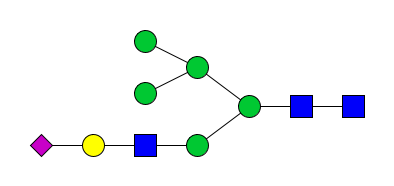 | H7N2 | 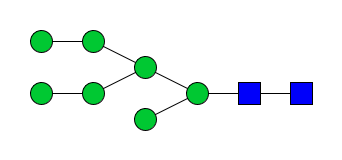 |
| H6N5F1S3 | 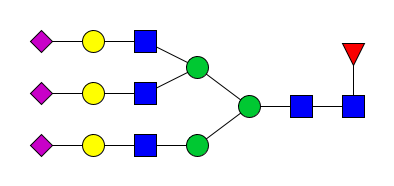 | H6N5S3 | 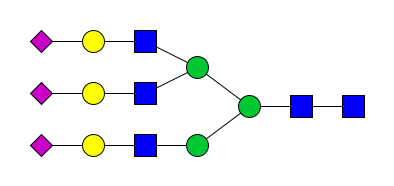 |
| H4N4F3S1 | 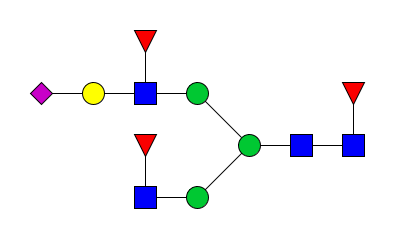 | H4N3 | 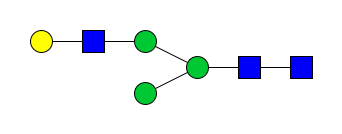 |
| H4N2 | 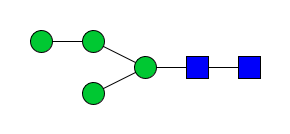 | H5N3S1 | 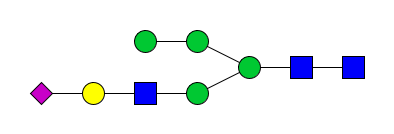 |
| H6N3F1S1 | 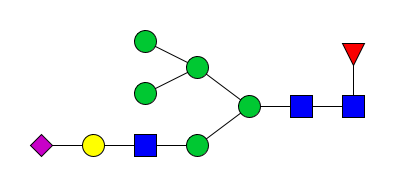 | H5N3F1S1 | 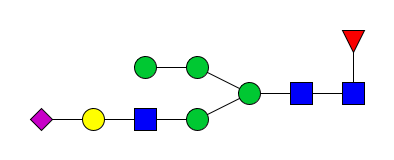 |
| H4N3F1S1 | 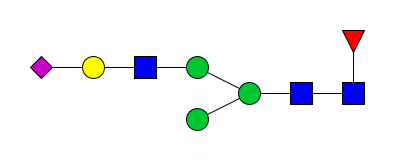 | H4N5F1S1 | 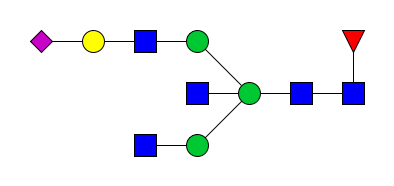 |
| H6N4S1 | 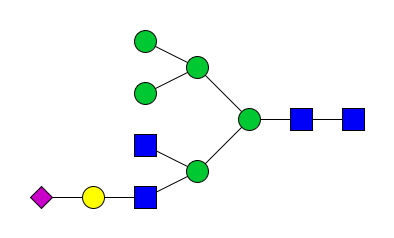 | H6N6 | 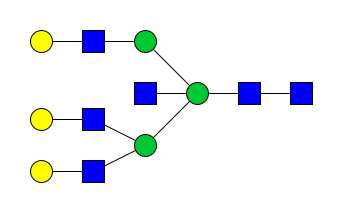 |
| H7N6S2 | 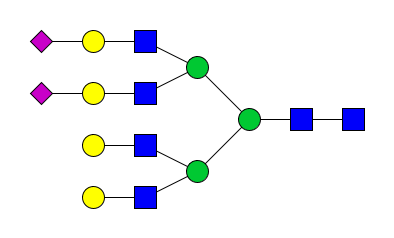 | H3N3 | 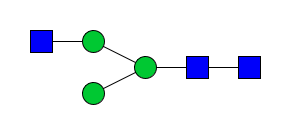 |
| H5N4F2 | 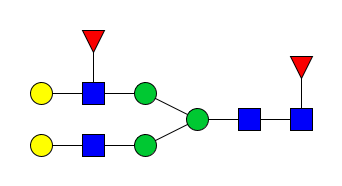 | H6N3 | 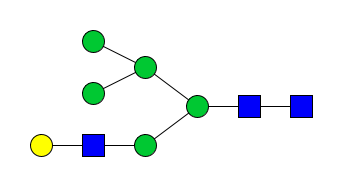 |
| H7N6S1 | 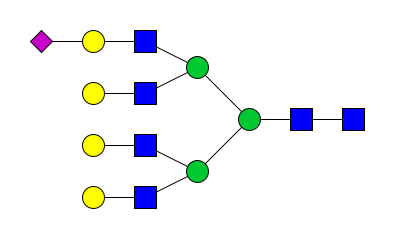 | H3N3F1 | 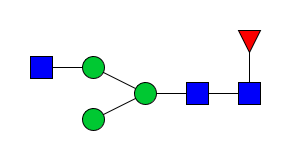 |
| H5N5S2 | 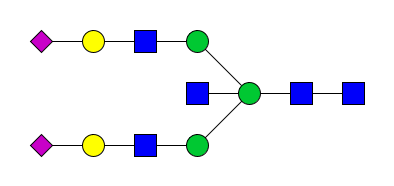 | H5N3 | 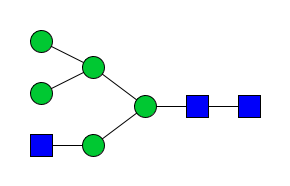 |
| H3N2F1 | 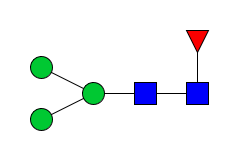 | H5N5F2 | 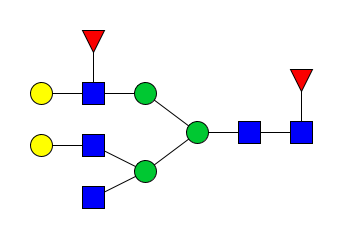 |
| H7N6S3 | 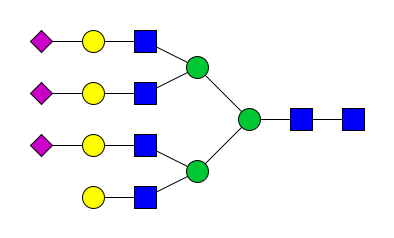 | H7N6F1S2 | 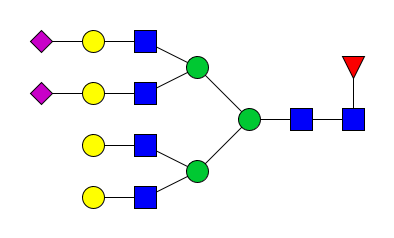 |
| H6N5F2S3 | 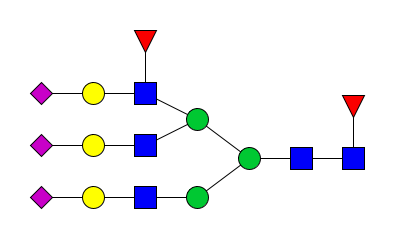 | H3N5F3 | 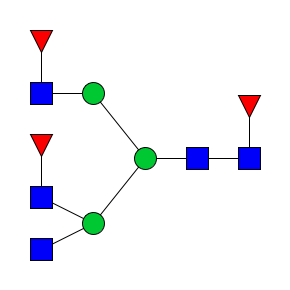 |
| H4N4F3 | 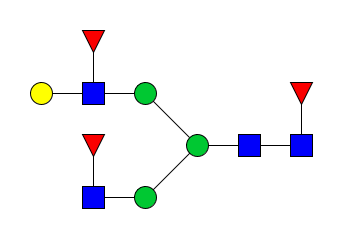 | H4N5S1 | 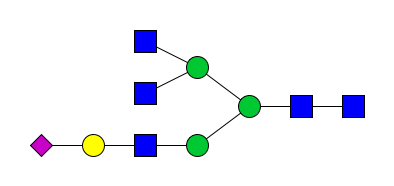 |
| H6N6F1S1 | 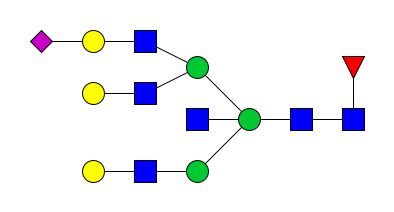 | H6N5F1 | 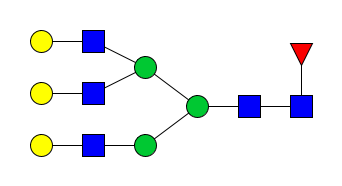 |
| H6N6F1S2 | 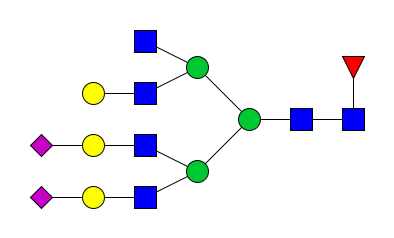 | H4N6S1 | 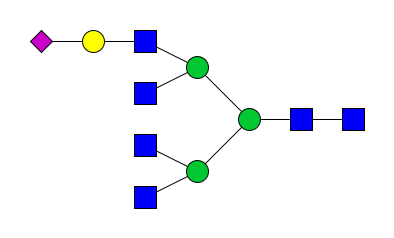 |
| H5N4F2S1 | 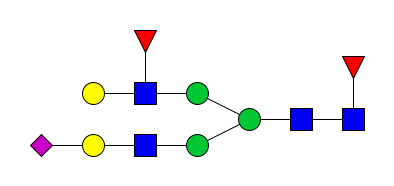 | H6N4S2 | 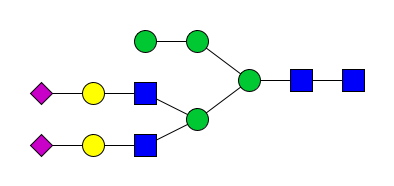 |
| H7N6F1S3 | 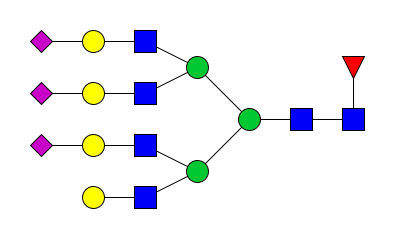 | H7N6F1S4 | 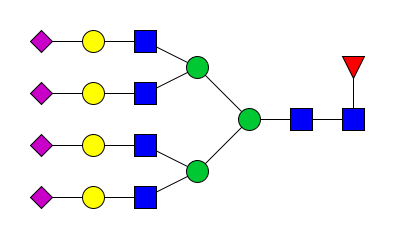 |
| H7N6S4 | 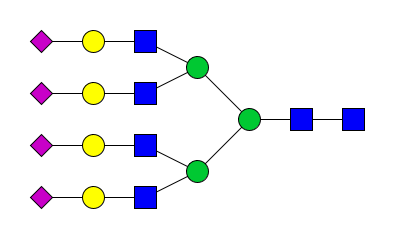 |  |  |

### Supplementary Note 1 – Instruction for writing custom GlyTrait formulas.

*The language style here is less formal, for the purpose is to teach the users of GlyTrait how to write their own formulas. For more details, and maybe the latest version of this instruction, please visit the Github page of GlyTrait:* <https://github.com/FudanLuLab/glytrait>.

The general format of a GlyTrait formula is:

**Name = [Numerator Terms] / [Denominator Terms]**

Let’s start with an example:

**CA1 = [(nAnt == 1) * (****type == ‘complex’)] / [****type == ‘complex’]**

Based on the general format, we can tell that the name of the trait is “CA1”, the numerator part is “(nAnt == 1) * (type == ‘complex’)”, and the denominator part is “type == ‘complex’”. The latter two parts seem a little confusing right now, which will be explained latter in detail. Putting it aside, the formula makes GlyTrait do the following calculation: “First find all glycans not only of complex type (type == ‘complex’), but also having one antenna (nAnt == 1), and calculate the total abundance of these glycans, denoted as $x_{num}$. Then calculate the total abundance of all complex glycans (type == ‘complex’), denoted as $x_{den}$. Finally, calculate $x_{num}/x_{den}$”. In short, this is the proportion of mono-antennary glycans within all complex glycans.

I hope this example gives you an intuition of how GlyTrait works. Let’s delve into the details of it. Basically, both the numerators and the denominators work in the same way: to calculate a weighted summation of the glycan abundances. Mathematically, what happens in the bracket [] could be generalized by:

$$x=\sum_{i}^{n} \theta_{i}a_{i}=\theta_{1}a_{1}+\theta_{2}a_{2}+\ldots+\theta_{n}a_{n}$$

where $n$ is the total number of glycans, and $a_{i}$ is the abundance of the i’th glycan. The weights ($\theta_{i}$) are determined by the content in []. For the denominator of CA1 (type == ‘complex’), $\theta_{i}$ is 1 if glycan $i$ is of complex type, 0 if not. Therefore, the resulting $x_{den}$ is the summation of the abundances for all complex glycans.

The numerator is more complicated. It contains two terms, one is (nAnt == 1), the other is (type == ‘complex’). You can probably guess out the meaning of the first term now. $\theta_{i}$ is 1 if glycan $i$ has only one antenna, 0 if not. The two terms are multiplied together with “*”. This results in an element-wise multiplication of the weights. If the weights of (nAnt == 1) are $\{\theta_{1}^{1}, \theta_{2}^{1}, \ldots, \theta_{n}^{1}\}$, and the weights of (type == ‘complex’) are $\{\theta_{1}^{2}, \theta_{2}^{2}, \ldots, \theta_{n}^{2}\}$, then the final weights are $\left\{ \theta_{1},\theta_{2}, \ldots, \theta_{n} \right\}=\{\theta_{1}^{1}\theta_{1}^{2}, \theta_{2}^{1}\theta_{2}^{2}, \ldots, \theta_{n}^{1}\theta_{n}^{2}\}$. For example, if glycan $i$ has more than one antenna, but it is a complex glycans, then $\theta_{i}^{1}=0$, and $\theta_{i}^{2}=1$, so $\theta_{i}=\theta_{i}^{1}\theta_{i}^{2}=0\times1=0$. In general, a numerator or denominator with more than one terms could be generalized by:

$$x=\sum_{i}^{n} \theta_{i}a_{i}=\sum_{i}^{n} ((\prod_{k}^{m} \theta_{i}^{k})a_{i})$$

where $m$ is the total number of terms. For example, if $m$ is 2 (like the numerator of CA1), it is:

$$x=\theta_{1}^{1}\theta_{1}^{2}a_{1}+\theta_{2}^{1}\theta_{2}^{2}a_{2}+\ldots+\theta_{n}^{1}\theta_{n}^{2}a_{n}$$

Now, let’s look at the details of the terms. What we have encountered, either (type == ‘complex’) or (nAnt == 1), are called the comparison terms. A comparison term compares a meta-property (a new terminology that will be explained latter) to a given value. Here, we used “==”, which means “is” or “equals to”. Note that there are two “=” instead of one, like the comparison operator in most programming language (you are reading this so probably you already know a bit about coding, but it’s worth mentioning anyway). Other comparison operators supported are “!=” (not equal to), “>” (greater than), “<” (less than”, “>=” (greater than or equal to), and “<=” (less than or equal to).

A meta-property is a conserved property automatically calculated by GlyTrait from glycan structure strings. There are three kinds of meta-properties: “categorical”, “numeric”, and “Boolean”. Supported meta-properties and possible values are:

| **Name** | **Definition** | **Type** | **Value Range** |
| --- | --- | --- | --- |
| type | the type of the glycan | categorical | “complex”, “hybrid”, or “high-mannose” |
| B | whether the glycan is bisected | Boolean | 1 or 0 |
| nAnt | the number of antennas | numeric | Integer ≥ 0 |
| nF | the number of fucoses | numeric | Integer ≥ 0 |
| nFc | the number of core fucoses | numeric | Integer ≥ 0 |
| nFa | the number of antennary fucoses | numeric | Integer ≥ 0 |
| nS | the number of sialic acids | numeric | Integer ≥ 0 |
| nM | the number of mannoses | numeric | Integer ≥ 0 |
| nG | the number of galactoses | numeric | Integer ≥ 0 |
| nN | the number of HexNAcs | numeric | Integer ≥ 0 |
| Pl | whether the glycan has poly-LacNAc | Boolean | 1 or 0 |
| nL | the number of α2,3-linked sialic acids | numeric | Integer ≥ 0 |
| nE | the number of α2,6-linked sialic acids | numeric | Integer ≥ 0 |

The categorical meta-properties only support “==” and “!=” operators. This is intuitive, as you can’t tell if a “type” is larger or smaller than something. A numeric meta-property supported all 6 operators. Finally, a Boolean meta-property can’t be used in a comparison term.

If you are familiar with coding, you will know that a comparison results in a Boolean value. That is, 0 or 1. So, (nAnt == 1) results in 0 or 1, and (type == ‘complex’) also results in 0 or 1. That’s why a Boolean meta-property can’t be used in a comparison term. If you want to “select bisected glycans with two antennas”, just use [B * (nAnt == 2)].

Besides the comparison terms, there are other two types of terms: the “direct terms”, and the “constant terms”. The direct terms are using meta-properties as terms directly. The “B” term in the last example is a direct term. Numeric meta-properties could also be used as a direct term. For example, in the formula of CFa (average number of antennary fucoses for complex glycans):

**CFa = [nFa * (type == 'complex')] / [type == 'complex']**

“nFa” here is a direct term. Let’s stop and think about what the formula means. The denominator is simple: “calculate the total abundance of complex glycans”. The numerator is more complex. As there are two terms, there are two weights for each glycan. $\theta_{i}^{1}$ is the number of antennary fucoses, and $\theta_{i}^{2}$ is whether the glycan is of complex type. The final weight of a glycan is $\theta_{i}=\theta_{i}^{1}\theta_{i}^{2}$. If the glycan is not of complex type, $\theta_{i}$ will be 0 anyway. If it is a complex type, $\theta_{i}$ will be the number of antennary fucoses of the glycan. So the numerator could be regarded as “the summation of complex glycan abundance weighted by number of antennary fucoses”.

If you are a little dizzy right now, let’s look at an example. Let’s say there are three glycans and only one sample. The compositions and relative abundance of the three glycans are:

| H5N4F1 | H5N2 | H5N4F2 |
| --- | --- | --- |
| 0.5 | 0.1 | 0.4 |

H5N4F1 and H5N4F2 are complex glycans, and only H5N4F2 has one antennary fucose. For CFa, the denominator is the total abundance of complex glycans: $x_{den}=1*0.5+0*0.1+1*0.4=0.9$. The numerator is: $x_{num}=0\left( no Fa \right)\times1\left( complex \right)\times0.5+0\left( no Fa \right)*1\left( not complex \right)\times0.1+1\left( one Fa \right)\times1\left( complex \right)\times0.4=0.4$. The final value of CFa for this sample is $0.4/0.9$. This is the average number of antennary fucoses for all complex glycans.

The last kind of term is much simpler, the constant term. It is just a number, more accurately, an integer. The most common use case of the constant terms is when you want to calculate “something per antenna”. For example, if you want to calculate the average sialic acids per antenna (or the average degree of sialylation for each antenna) within bi-antennary glycans, there we have

**A2S = [(nS / 2) * (type == 'complex') * (nAnt == 2)] / [****(type == 'complex') * (nAnt == 2)]**

Note that the direct term “nS” is divided by a constant term “2”. “/” here is another linkage operator for terms, other than “*”. Note that “/” could only be used before a numeric direct term, or a constant term. With “/”, you could calculate something like “average degree of sialylation for each galactose”, as in

**A2GS = [(nS / nG) * (type == 'complex') * (nAnt == 2)] / [****(type == 'complex') * (nAnt == 2)]**

You might have noticed that there was a lot of typing in the formulas. Like in the formula of A2GS, we had to type “(type == 'complex') * (nAnt == 2)” twice for the numerator and the denominator. Luckily, GlyTrait supports a shortcut for this. Just use [] // [] instead of [] / [], GlyTrait will add all terms in the denominator to the numerators. The formulas before looks more concise right now:

**CA1 = [nAnt == 1] // [type == ‘complex’]**

**CFa = [nFa] // [type == ‘complex’]**

**A2S = [nS / 2] // [(type == 'complex') * (nAnt == 2)]**

**A2GS = [nS / nG] //** **[(type == 'complex') * (nAnt == 2)]**

This is the default format for all GlyTrait built-in formulas (with some exceptions not able to be expressed in this way). This shortcut grammar also makes the formula more comprehensible. It allows a general interpretation of “something within some type of glycans”. For example, “average number of sialic acids per galactose (nS / nG) within all bi-antennary (nAnt == 2) complex (type == ‘complex’) glycans” for A2GS. Note that in this case, all terms in the denominator should be comparison terms or Boolean direct terms.

Finally, the parentheses () are just for visual separation of terms in the formulas. GlyTrait will just ignore them when parsing the formulas. However, for explicitness and less confusion, we recommend using them around comparison terms, except it is the only one in the denominator or the numerator.

That’s all you need to know about how to write a formula. For a reference, look at the built-in formulas in Supplementary Table 1. Happy using GlyTrait!

### Supplementary Note 2 - The derived trait calculation algorithm of GlyTrait

The calculation of derived traits consists of three steps.

In the first step, input files are loaded and parsed. The glycan abundance table is pre-processed according to the procedure described above, stored as a DataFrame in pandas package (v2.0.1). The glycoCT formatted glycan structures are parsed by the glypy package (v1.0.11) and stored as tree structures. The built-in and user-provided formula files are combined and parsed by the built-in parsing module.

In the second step, a meta-property table is generated. A meta-property of a glycan is either a Boolean value, an integer, or a category variable, reflecting a basic property of glycan structures. All meta-properties considered by GlyTrait include “glycan type”, “bisection”, “number of antennas”, “number of core fucoses”, “number of antennary fucoses”, “number of sialic acids”, “number of mannoses”, “number of galactose”, “number of GlcNAc”, “whether the glycan has poly-LacNAc”, “number of α2,3-linked sialic acids”, “number of α2,6-linked sialic acids”. Tree structures of each glycan is traversed in either depth-first or breath-first manner to acquire meta-property values. Due to the difficulties of linkage determinations of N-glycans with regular mass spectrometry, GlyTrait support handling of structural ambiguity. The resulting meta-property table $D_{mp}$ is a DataFrame with glycans as indexes and meta-property names as columns.

In the final step, the meta-property table is used as reference to calculate derived traits in a vectorized manner. The formulas are first initialized on the meta-property table to get two coefficient vectors, one for the numerator ($\boldsymbol{C}^{n}$) and the other for the denominator ($\boldsymbol{C}^{d}$). The coefficients are the weights of each glycan, differed by the definition of the derived trait. Please see Supplementary Note 1 for information about formula design and coefficient calculation. Assume that N is the total number of glycans, M is the total number of samples, O is the total number of derived traits. Derived trait $j$ for sample $k$ is calculated by

$$t_{j,k}=\frac{\boldsymbol{C}_{j}^{n}\cdot\boldsymbol{A}_{k}}{\boldsymbol{C}_{j}^{d}\cdot\boldsymbol{A}_{k}}=\frac{\sum_{i}^{N} c_{i, j}^{n}a_{i}}{\sum_{i}^{N} c_{i, j}^{d}a_{i}}$$

where $\boldsymbol{C}_{j}^{n}=\left( c_{1, j}^{n}, c_{2, j}^{n}, \ldots, c_{i, j}^{n}, \ldots,c_{N, j}^{n} \right)$ is the numerator coefficient vector for derived trait $j$, $\boldsymbol{C}_{j}^{d}=\left( c_{1, j}^{d}, c_{2, j}^{d}, \ldots, c_{i, j}^{d}, \ldots,c_{N, j}^{d} \right)$ is the denominator coefficient vector for derived trait $j$, $\boldsymbol{A}_{k}=\left( a_{1},a_{2}, \ldots,a_{i},\ldots,a_{N} \right)$ is the glycan abundance vector of sample $k$. The actual computation of derived traits leverages numpy (v1.25.2) and pandas (v2.0.1) package for vectorization, obtaining a derived trait vector $\boldsymbol{T}_{j}$ for all samples. Finally, all derived trait vectors $\left\{ \boldsymbol{T}_{1},\boldsymbol{T}_{2}, \ldots,\boldsymbol{T}_{j},\ldots,\boldsymbol{T}_{O} \right\}$ are concatenated into a DataFrame, with sample names as indexes and derived trait names as columns.

### Supplementary Note 3 – The post-filtering algorithm of GlyTrait

The post-filtering consists of two steps: the low-variance filtering step and the collinearity filtering step. The former removes derived traits with low variance across samples, and the later prunes the derived trait dimension to reduce information redundancy.

The low-variance filtering starts with removing derived trait with missing values appear in more than half the samples. Missing values could be induced when a derived trait could not be calculated for a certain glycan set. For example, if a glycan set has no tetra-antennary glycans, A4Fc (the proportion of glycans with core fucosylation within all tetra-antennary glycans) will be missing values in all samples. After removing highly missing derived traits, those with zero variance across all samples are also removed. These normally include derived traits that provides no useful information to the research questions. For example, the CHO EPO has no glycans with bisection, so all derived traits about bisection (e.g. CB, A2B) will be zero.

The collinearity filtering starts with building two networks with derived traits as nodes, one correlation network from the calculated derived traits after low-variance filtering, and one ontology network from the definition of the derived traits. In the correlation network, an edge exists between two derived traits if the correlation coefficient exceeds a predefined threshold. Mathematically, let $D=\left\{ d_{1},d_{2},\ldots,d_{O} \right\}$ be the set of derived traits after low-variance filtering, then the correlation network could be represented as $G_{c}=\left( V, E_{c} \right)$, where $V$ is the set of nodes corresponding to the derived traits in $D$, and $E_{c}$ is the set of edges between nodes. An edge $\left( d_{i}, d_{j} \right)\in E_{c}$ if and only if Pearson’s correlation coefficient $corr\left( d_{i}, d_{j} \right)>\theta$, where $\theta$ is the predefined threshold. The ontology network could be represented as $G_{O}=\left( V, E_{O} \right)$, with similar notations as the correlation network. However, $G_{O}$ is a directed acyclic graph (DAG), in which any edge in the network is directed. Briefly, a directed edge $\left( d_{i}, d_{j} \right)\in E_{O}$ exists if and only if $d_{i}$ is “one step more general” than $d_{j}$. For example, an edge exists from “A2Fc” to “A2SFc”, and another edge exists from “A4B” to “A4FB”. The concrete criteria for the edges in $E_{O}$ are explained in Supplementary Note 4.

The two networks are then merged into a single network, such that an edge between two nodes exists if and only if it exists in both the correlation and ontology networks, retained the directed nature of the ontology network. Mathematically, the merged network could be represented as $G_{M}=\left( V, E_{M} \right)$, where $V$ is the same as $G_{c}$ and $G_{O}$, and $E_{M}=E_{c}\cap E_{O}$. Computationally, the merged network is calculated by element-wise logical AND operating for the adjacency matrices from the two original networks. Finally, only derived traits with zero in-degree in the merged network are retained. This process ensures that filtering will only be activated when two derived traits with both semantic and data-wise similarity, retaining potentially meaningful correlation relationships. Network analysis outside the GlyTrait framework was performed with Python NetworkX library (v2.8.0), while the calculation inside GlyTrait is directly based on Python numpy library (v1.25.2).

### Supplementary Note 4 – Building the derived trait ontology network

To perform the collinearity filtering (see Methods), an ontology network of derived traits has to be generated. One option was to integrate a manually curated hard-coded network (Supplementary Fig. 4) inside GlyTrait. However, as GlyTrait support custom formulas, we designed an algorithm to build the network dynamically.

In GlyTrait, the formulas are parsed into a list of numerator terms and a list of denominator terms (see Supplementary Note 1 for the design of GlyTrait formulas). Specially, a term after the division operator “/” is wrapped into a new term. For example, for the formula of A2S:

**A2S = [nS / 2] // [(****type == 'complex') * (****nAnt == 2)]**

the numerator terms are: “nS”, “/ 2”, “type == 'complex'”, and “nAnt == 2”; the denominator terms are: “type == 'complex'” and “nAnt == 2”. Note that when “//” is used, the denominator terms will be added to the numerator terms.

Assume the ontology network is $G_{O}=\left( V, E_{O} \right)$, and $D=\left\{ d_{1},d_{2},\ldots,d_{O} \right\}$ is the set of derived traits after low-variance filtering. A directed edge $\left( d_{i}, d_{j} \right)\in E_{O}$ exists if both the numerator and the denominator of $d_{j}$ include an additional term compared to $d_{i}$, which must be the same term.

For example, see the formula of A2FS:

**A2FS = [nS / 2] // [(type == 'complex') * (nAnt == 2) * (nF > 0)]**

Again, “//” is used here, so the new term (nF > 0) will be added both to the numerator and the denominator, compared to A2S. As a result, A2FS should be a “child” of A2S in the ontology network.

### Supplementary Figure 1 – Computational framework of GlyTrait

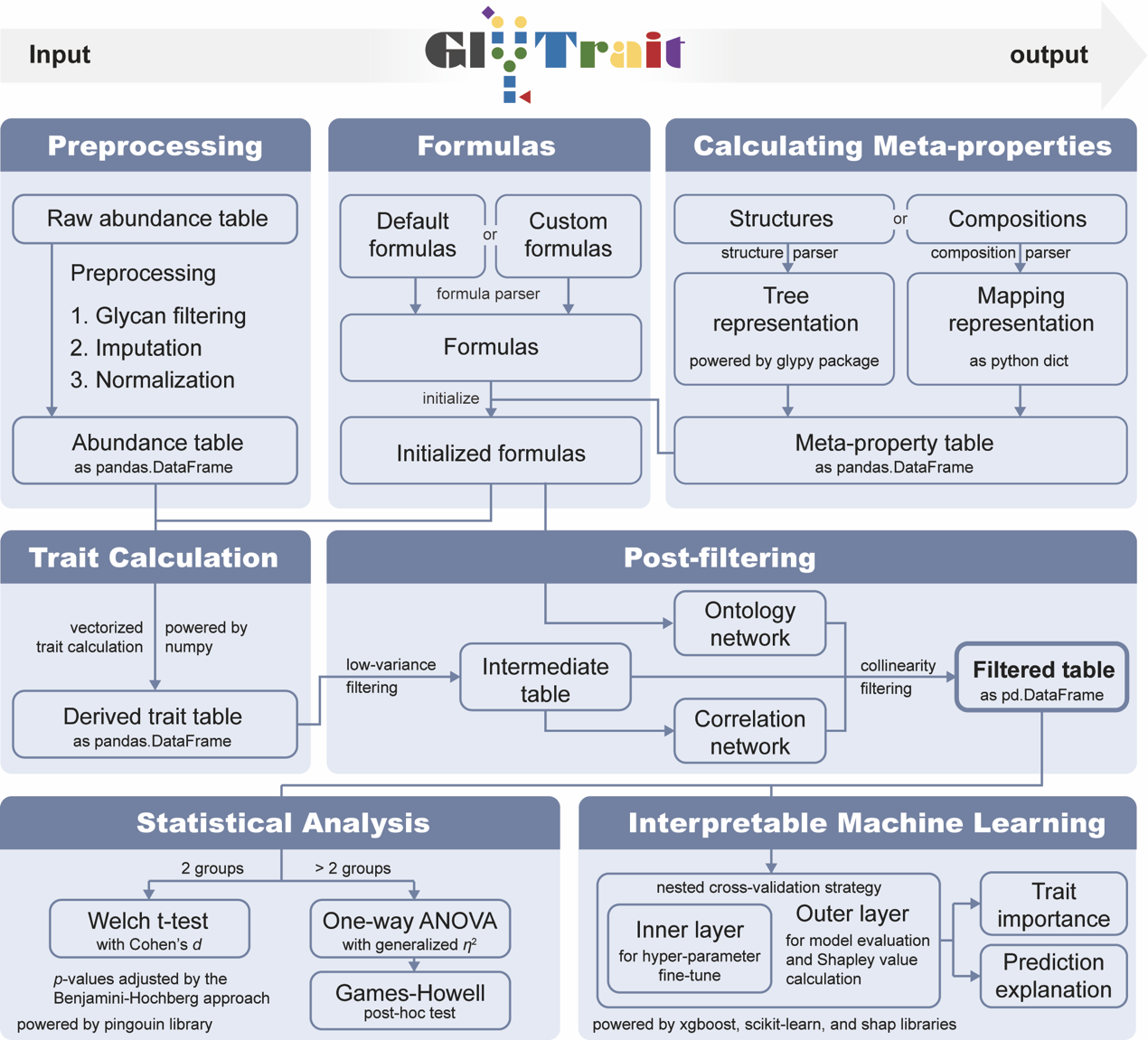

### Supplementary Figure 2 – Derived traits in the CHO dataset

This heatmap shows the values of all derived traits after post-filtering with a correlation threshold of 1.0 for the CHO dataset. Grey tiles are missing values. This could result from missing certain glycans in certain samples. For example, all traits starting with “A4” are missing for KO:mgat5, for tetra-antennary glycans are not expressed in this sample.

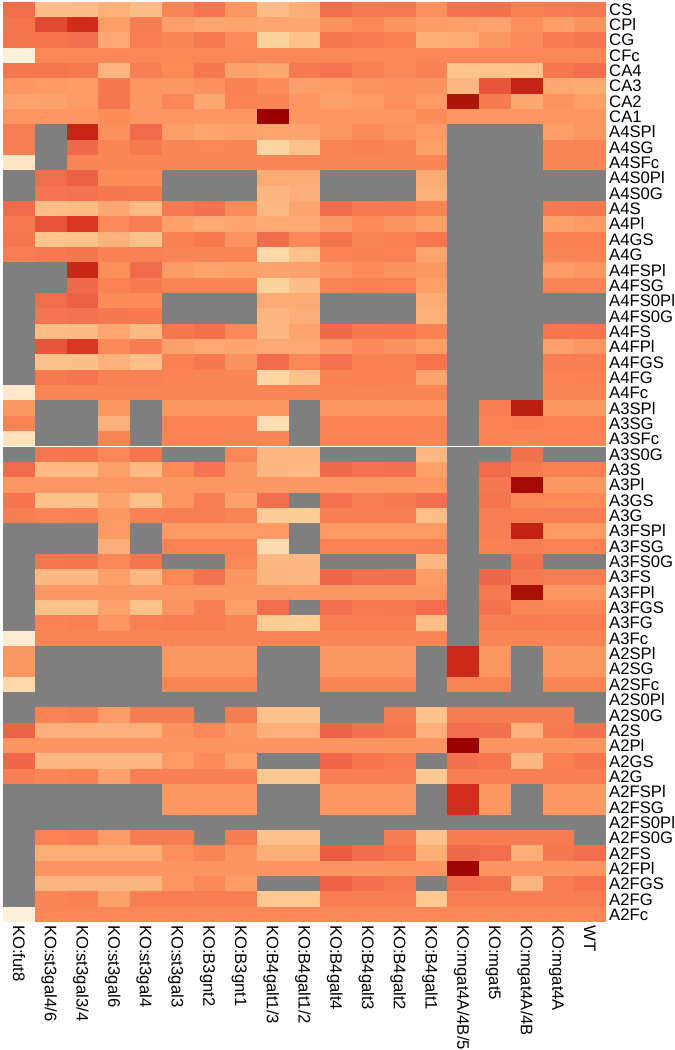

### Supplementary Figure 3 – Correlation for derived traits in the CHO dataset

The correlation heatmap of all derived trait after post-filtering with a correlation threshold of 1.0 for the CHO dataset. Grey tiles mean one of the traits compared contains too many missing values so the correlation analysis is not available (see the R corrplot package v0.92 for details).

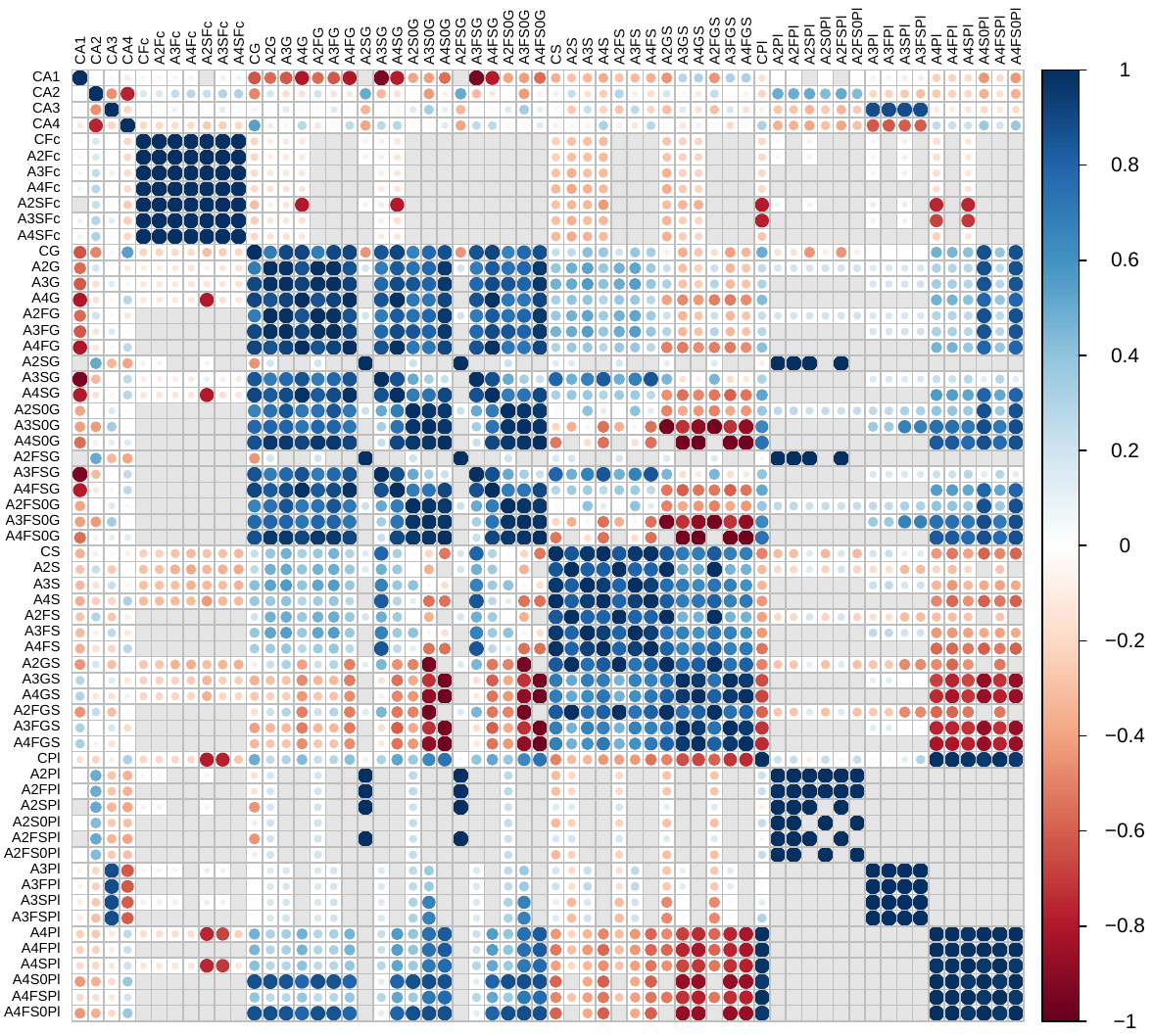

### Supplementary Figure 4 – Trait ontology network (partial)

The trait ontology network without sialic acid linkage traits and traits about poly-LacNAc. Purple: traits about sialylation. Yellow: traits about galactosylation. Blue: traits about bisection. Red: traits about fucosylation. Brown: global traits without children or parents.

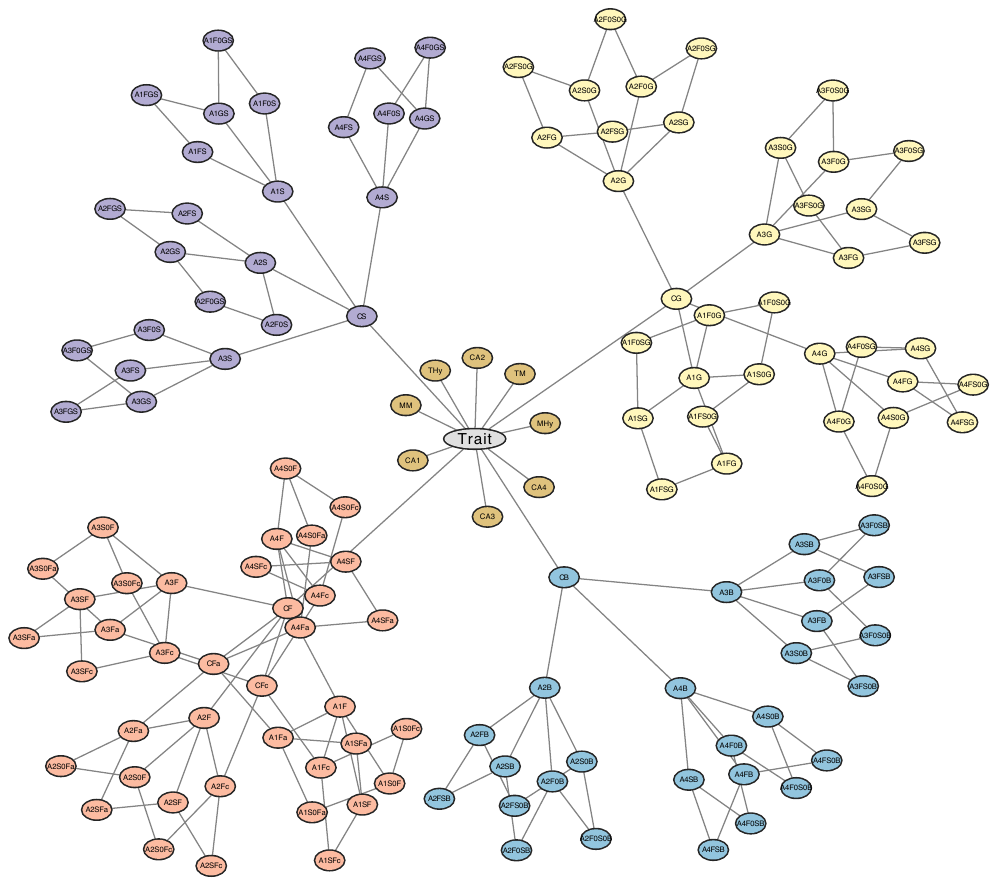

### Supplementary Figure 5 – Trait ontology network (full)

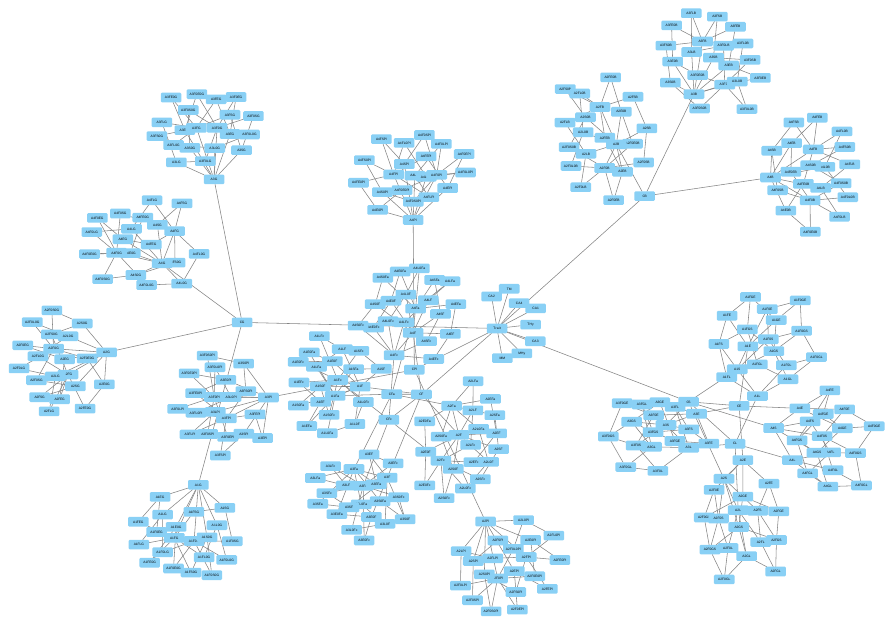

The trait ontology network containing all built-in structural derived traits.

### Supplementary Figure 6 – Information loss of collinearity filtering

Machine learning model comparison for the 65 derived traits kept by post-filtering with correlation threshold setting to 0.9. Accuracy, ROC AUC, F1 scores and PR ROC were acquired by a repeated 5-fold cross validation (repeated 10 times). All p-values were acquired by Wilcox rank sum test with Benjamini-Hochberg multiple comparison adjustment. CART: Classification and Regression Tree, LR: Logistic Regression, NB: Naïve Bayes, RF: Random Forest, SVM: Support Vector Machine. ns: not significant, *: 0.01 < p < 0.05, **: 0.001 < p < 0.01, ***: p < 0.001.

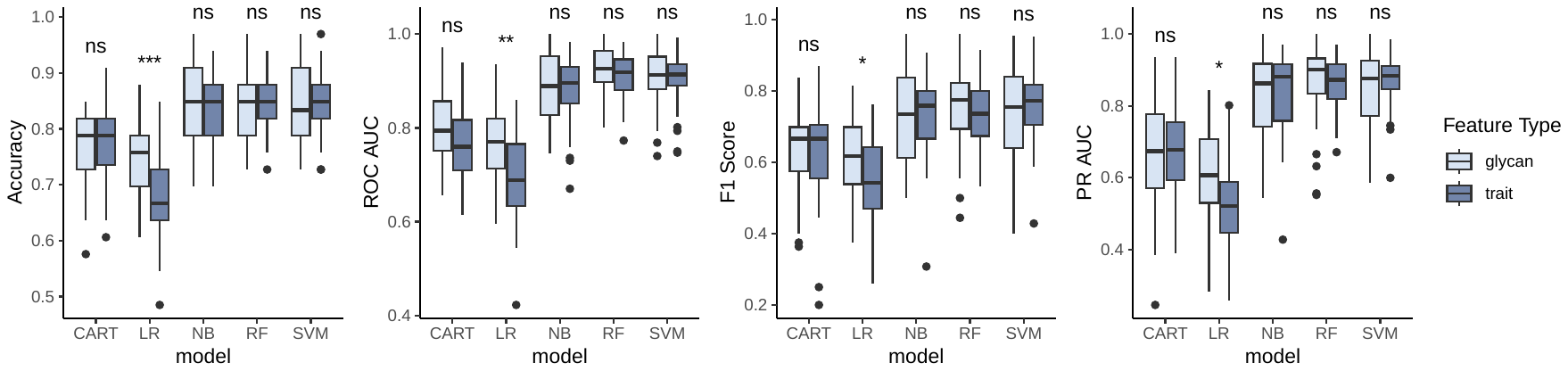

### Supplementary Figure 7 – Statistical analysis for CFc on simulated data

The boxplots of CFc (proportion of core-fucosylated glycans within all complex glycans) for baseline samples and each fold change condition. The p-values were acquired from Welch t-test adjusted by Benjamini-Hochberg method. Each box contains 100 simulated samples (see Methods).

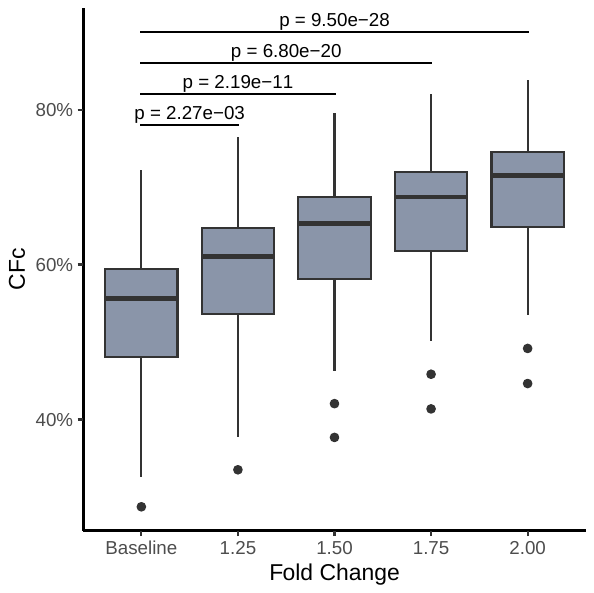

### Supplementary Figure 8 – Statistical analysis for glycans on simulated data

The distributions of FDR for glycans in Welch t-test between the baseline samples and each fold change conditions. There were 78 glycans in total, and 0, 3, 74, 78 glycans show statistical significance (FDR < 0.05) between baseline samples and samples with fold change of 1.25, 1.50, 1.75, and 2.00.

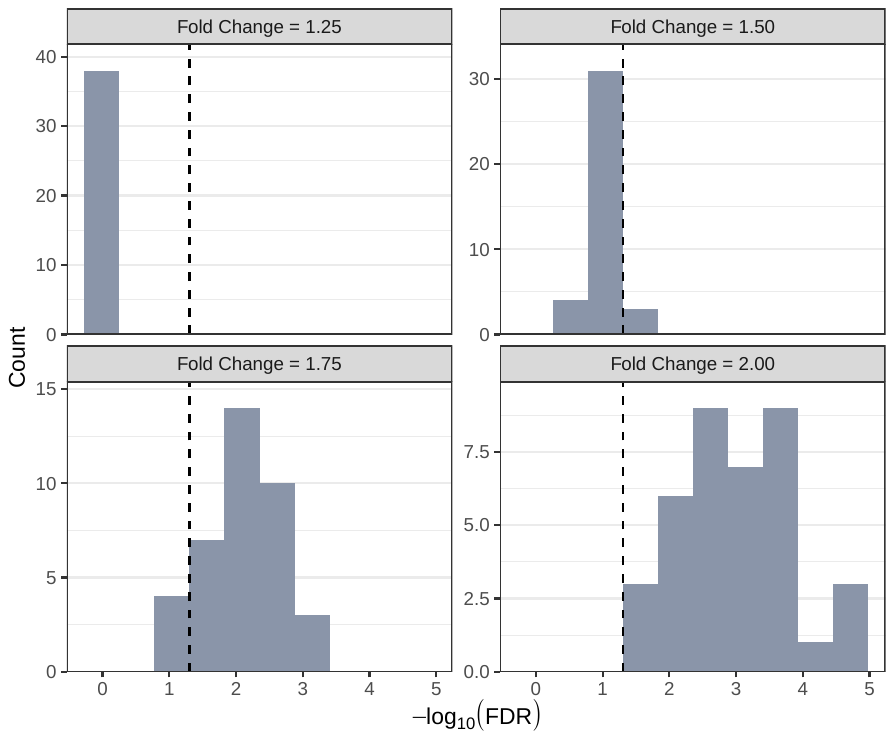

### Supplementary Figure 9 – Boxplots for the 36 significant derived traits in HCC dataset

All p-values are from Welch t-test with Benjamini-Hochberg adjustment.

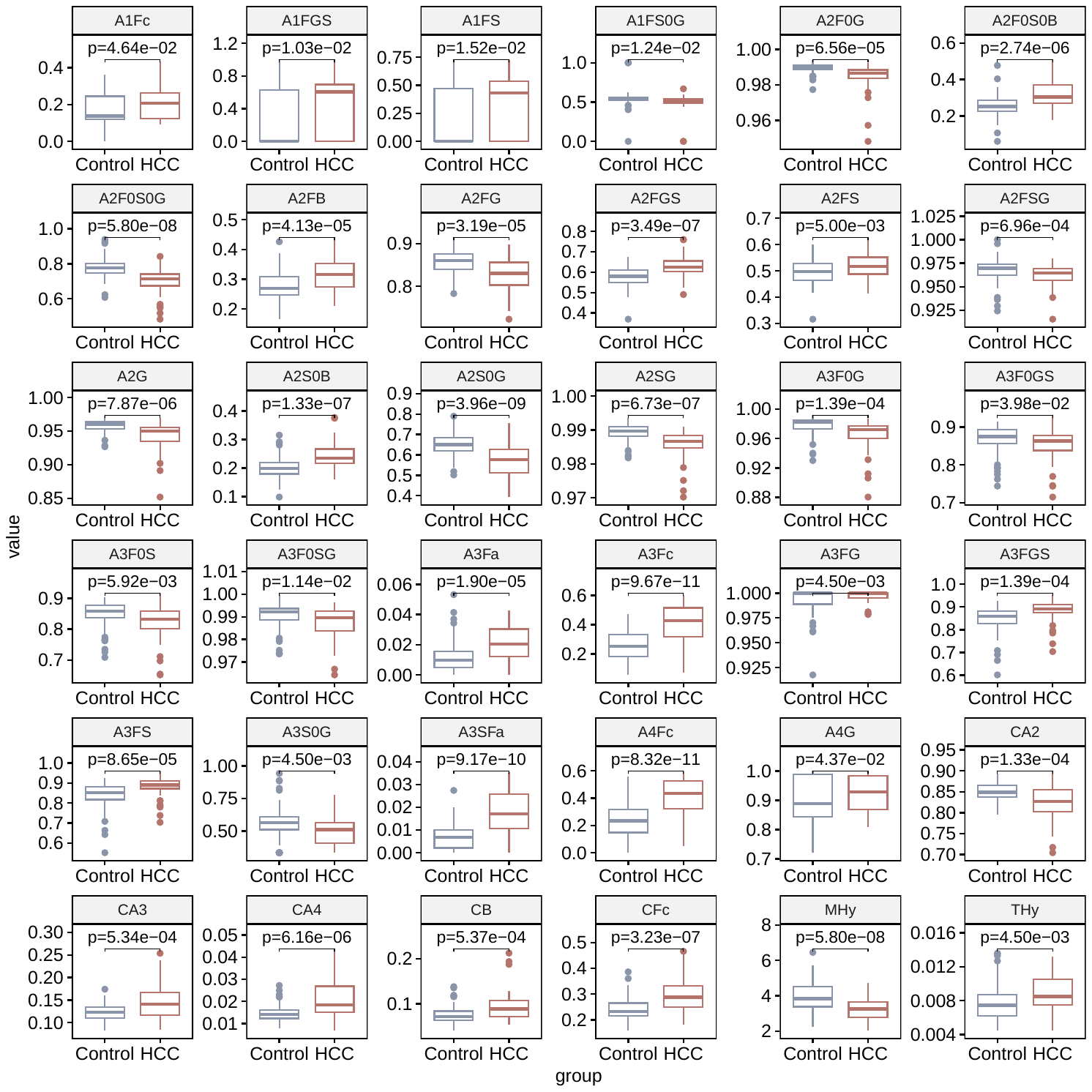

### Supplementary Figure 10 – ROC curves of derived trait in HCC dataset

This figure includes the ROC curves of all derived traits with a lower bound of the 95% CI for ROC AUC exceeding 0.6. The grey lines are the raw ROC curves. The light blue lines are the ROC curves from 100 bootstraps. The 95% CIs are calculated from 1000 bootstraps.

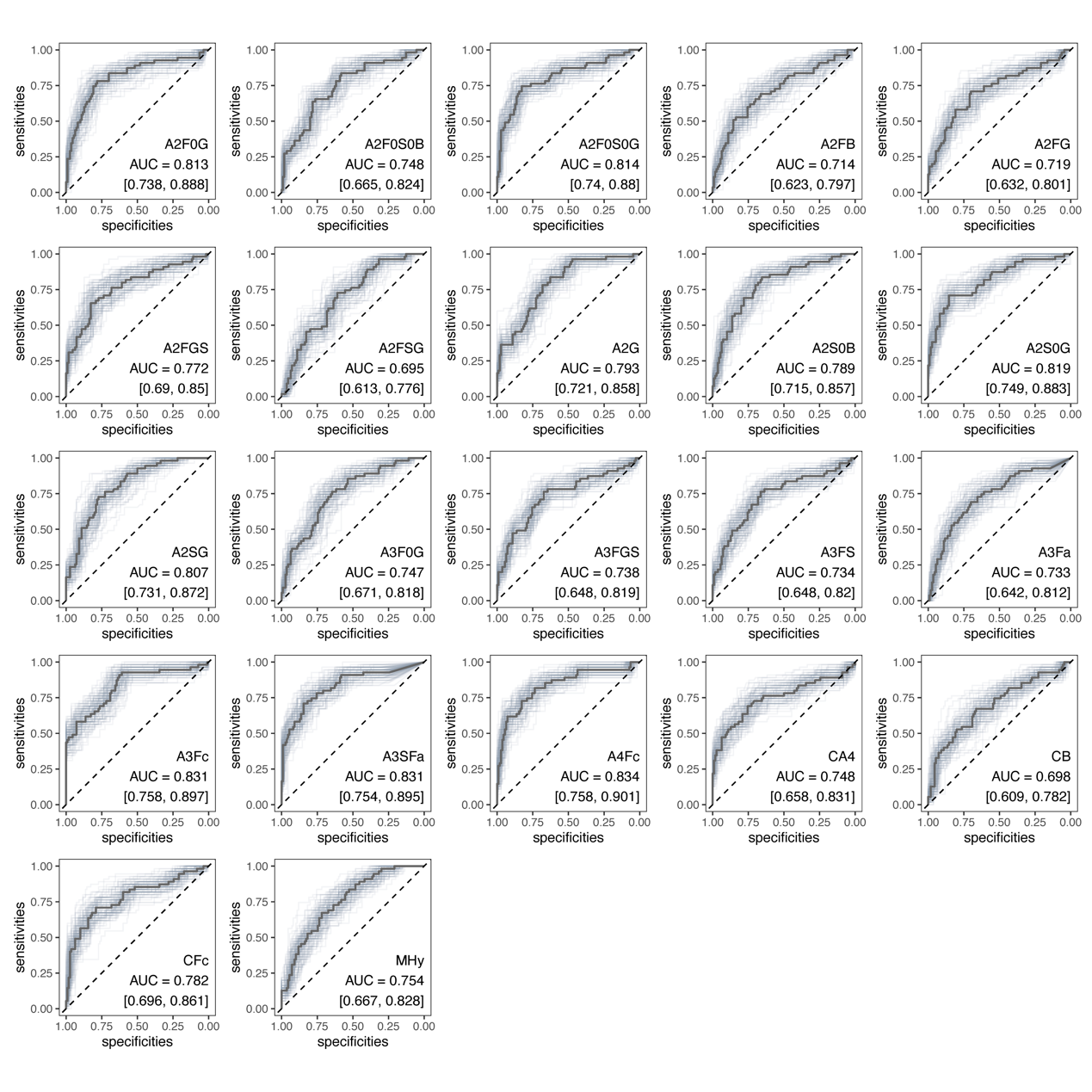

### Supplementary Figure 11 – Nested cross-validation strategy

The cross-validated strategy for hyperparameter fine-tune, model evaluation, and Shapley value calculation. The inner layer ensures a well-performed model. The outer layer ensures Shapley value calculation for all available samples, and an unbiased evaluation of the model.

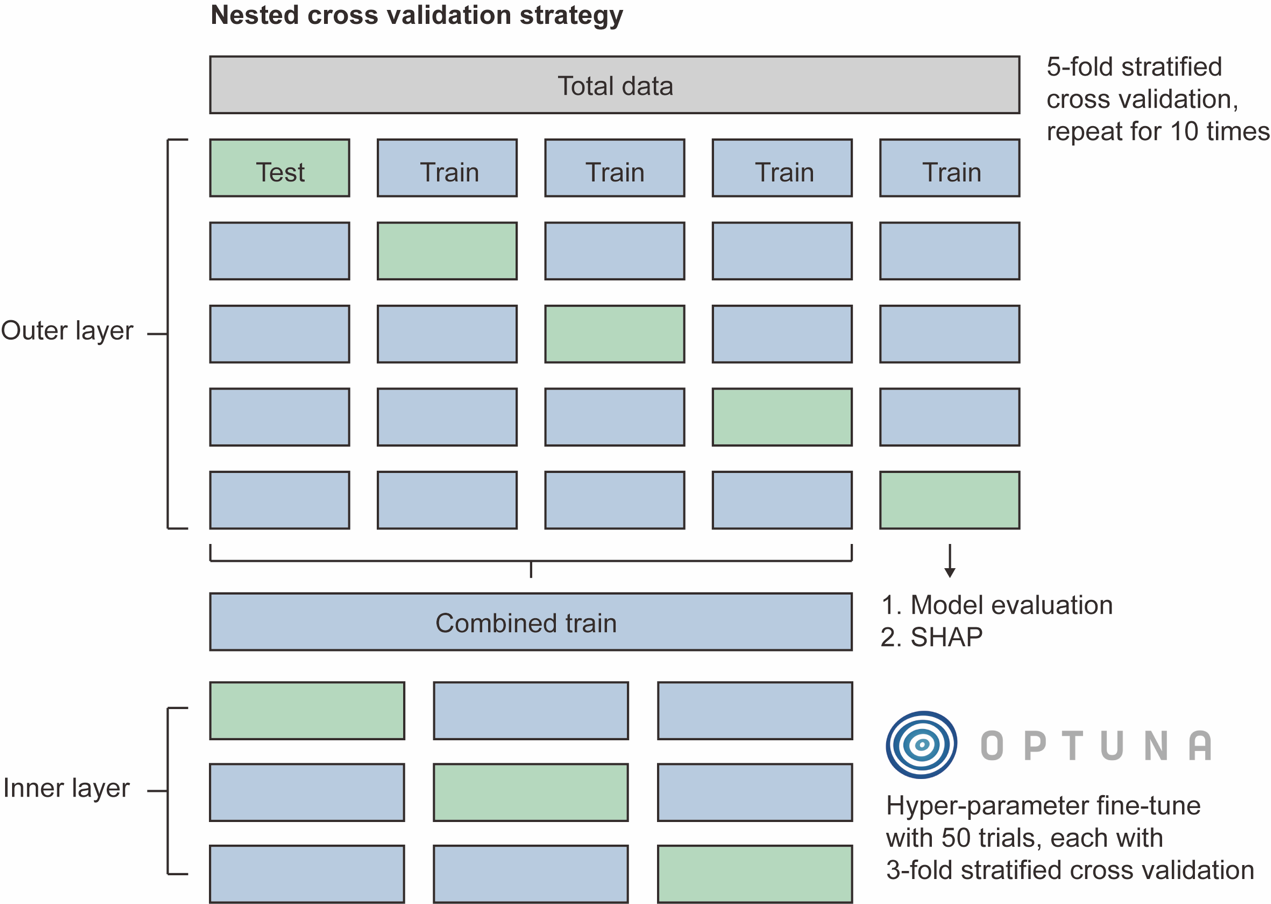
